# BinSPreader: refine binning results for fuller MAG reconstruction

**DOI:** 10.1101/2022.02.14.480326

**Authors:** Ivan Tolstoganov, Yuri Kamenev, Roman Kruglikov, Sofia Ochkalova, Anton Korobeynikov

## Abstract

Despite the recent advances in high-throughput sequencing, analysis of the metagenome of the whole microbial population still remains a challenge. In particular, the metagenome-assembled genomes (MAGs) are often fragmented due to interspecies repeats, uneven coverage and vastly different strain abundance. MAGs are usually constructed via a dedicated binning process that uses different features of input data in order to cluster contigs that might belong to the same species. This process has some limitations and therefore binners usually discard input contigs that are shorter than several kilobases. Therefore, binning of even simple metagenome assemblies can miss a decent fraction of contigs and resulting MAGs oftentimes do not contain important conservative sequences that might be of great interest of researcher.

In this work we present BinSPreader — a novel binning refiner tool that exploits the assembly graph topology and other connectivity information to refine the existing binning, correct binning errors, propagate binning from longer contigs to shorter contigs and infer contigs belonging to multiple bins. Furthermore, BinSPreader can split input reads in accordance with the resulting binning, predicting reads potentially belonging to multiple MAGs. We show that BinSPreader could effectively complete the binning, increasing the completeness of the bins without sacrificing the purity and could predict contigs belonging to several MAGs.

## 1 Introduction

Amount of microbial organisms which can be easily cultivated is relatively small in proportion to the Earth’s total diversity [28], therefore most of the Earth’s microbiota proves difficult for analysis. Whole metagenomic shotgun sequencing, which allows for a comprehensive analysis of microbial DNA from a sample, provides an alternative method of understanding of the functional potential and genetic composition of different microorganisms that have not been previously cultured. Metagenomic sequencing libraries are then assembled using metagenomic assemblers, such as metaSPAdes [26] or MEGAHIT [12] for short read libraries, or metaFlye [11] for long read libraries.

In order to extract useful information from complex metagenomic assemblies, a process called *binning* is used. State-of-the-art binners use all different kinds of information including nucleotide content, observed contig abundance, paired-end read connectivity and other connectivity (e.g. from Hi-C links [5]) to cluster contigs that might belong to the same species. However, this kind of information could only be considered reliable for long contigs and therefore the majority of binners discard contigs that are shorter than several kilobases. Nevertheless, the set of contigs could not be considered as the ultimate result of a metagenomic assembly. Indeed, the complete information about the assembly is provided via the assembly graph. Usually the edges of an assembly graph are the maximal non-branching genomic sequences (unitigs) and the resulting contigs are paths in this assembly graph obtained after the repeat resolution process. Recent development of such assembly graph-aware alignment tools like SPAligner [7], PathRacer [34], GraphAligner [29] among the others shows that the proper utilization of the assembly graph could significantly improve the obtained results.

To date, it seems that the connectivity information between the contigs in the assembly graph is ignored by the majority of the common binning tools like MetaBAT2 [10], MetaWrap [37], and VAMB [25] potentially reducing the overall precision of the results. Recently developed graph-aware binning refining tools such as METAMVGL [39], MetaCoAG [15] and Binnacle [21] also do not utilize the assembly graph in the usual sense of the term. Instead they are relying on the so-called *scaffold graph* that only preserves the connectivity information between different scaffolds. However, the original assembly graph contains more information including the multiplicity of edges and the set of edges that comprise a contig. In order to utilize this greater amount of information we suggest using the original assembly graph instead of scaffold graph. This brings to us many opportunities such as multiple binning of individual edges, binning correction and more precise bin label propagation (from edge to edge and not from scaffold to scaffold).

Standard MAG quality assessment tools, such as AMBER [18] and CheckM [27] do not assess MAGs for the presence of important sequences, such as mobile genetic elements (MGEs), antibiotic resistance genes (AMR) and CRISPR arrays, that have very high agricultural or clinical importance. As such, MAGs with over 80% completeness as reported by AMBER or CheckM may contain less than 45% of genomic islands and less than 30% of plasmid sequences [14]. Mobile genetic elements are commonly flanked by direct repeats [30], and are therefore located on short repetitive edges of the assembly graph and associated with multiple organisms.

Besides MGEs, MAGs often miss contigs containing rRNA genes. Bacterial genomes contain multiple copies of ribosomal genes forming tangled repeat structures which are often not assembled well. In a metagenome the situation is further complicated by the presence of conservative parts of rRNA genes shared between different species. Such sequences form intra- and interspecies repeats and therefore the overall recovery of a decent-length rRNA genes sequences from a metagenome assembly is quite low [17]. Finally, the contigs containing rRNA genes have different abundance (due to high copy number) and nucleotide content effectively preventing the majority of binning attempts. As such, inclusion of short edges of the assembly graph into MAGs is crucial for detecting MGE and rRNA sequences.

In this work we show that assembly graph representation provides more accurate multiple binning of short edges that scaffold graph representation. We present a new software tool BinSPreader which can produce refined MAGs from initial binning by combining metagenomic assembly graph and sequencing data. We show that BinSPreader can accurately predict contigs belonging to multiple bins and besides improving the usual completeness / purity metrics of MAGs is able to recover MGE and rRNA sequences more accurately than state-of-the-art binning refining tools. BinSPreader is available from cab.spbu.ru/software/binspreader.

## 2 Results

### 2.1 Datasets

We used several mock metagenomic datasets, simulated metagenomes as well as real metagenomes for the refining evaluation. These metagenomes are derived from different communities exhibiting different microbial composition, abundance profiles, genome characteristics and similarity intended to provide a broader scope of binning data features.

**MBARC26** [36] is composed of 23 bacterial and 3 archaeal strains isolated from heterogeneous soil, aquatic environments as well as human, bovine and frog microbiota. The genomes of these species span a wide range of genome sizes (1.8–6.5 Mbp), GC-contents (28.4–72.7%) and repeat contents (0–18.3%).

**BMock12** [32] includes DNA from 12 bacterial strains belonging to actinobacterial, flavobacterial and proteobacterial taxa that also display a large spread of genome properties. Apart from this, it includes three bacteria with genomes of high %GC and average nucleotide identity (ANI) which complicates the assembly and binning.

ZymoBIOMICS Microbial Community Standard (referred as **Zymo**) is a mock community consisting of eight bacterial and two fungal strains. These organisms are lysed in varying degrees and significantly differ in terms of the completeness of sample DNA extraction, which is a determining factor for sequencing and downstream analysis.

The benchmarking dataset from [14] (referred as **magsim-MGE**) contains paired-end Illumina sequencing data of 30 bacteria with randomly assigned relative abundance. It is designed to display a high diversity of genetic features, such as plasmids and genomic islands.

We assembled each of these datasets from Illumina shotgun sequencing using metaSPAdes 3.15.3 and used reference genomes of included bacteria, archaea and yeasts to construct ground truth binning standards for benchmark studies.

**simHC+** simulated dataset [38] was derived out of genome assemblies of 100 bacterial species that mimics high-complexity communities lacking dominant strains. As no original reads for this dataset was available, we used metagenomic assembly, abundance profiles and ground truth binning standard as provided in MetaCoAG paper [15].

**IC9** is a real clinical gut metagenome of a chronically critically ill patient collected in a critical care unit. The dataset contains both paired-end and Hi-C data which was crucial for better resolution of MAGs [8]. The metagenome is harboring many antibiotic-resistant strains with elevated levels of horizontal gene transfer. The dataset was assembled as described in [8].

**Sharon** dataset [33] contains the metagenomic sequencing data of pre-born infant fecal samples collected across 18 time points. All these sequencing libraries were co-assembled together using metaSPAdes 3.15.3 before binning and refining.

### 2.2 Evaluated approaches

We benchmarked BinSPreader against state-of-the-art graph-aware binning refiners METAMVGL [39], MetaCoAG [15] and Binnacle [21], as well as consensus-based refiner DAS_TOOL [35]. While all five binning refiners require metagenomic assembly, their requirements for other types of input data differ.

MetaCoAG, Binnacle, and BinSPreader require assembly graph in GFA format as an input. METAMVGL utilizes assembly graph in obsolete FASTG format which makes it impossible to use on assembly graphs produced by e.g. metaFlye. METAMVGL, Binnacle, DAS_TOOL and BinSPreader require initial binning to refine, while MetaCoAG produces initial binning internally using provided coverage profiles. Paired-end read library is required for both METAMVGL and Binnacle as a source of connectivity information between scaffolds and for BinSPreader input paired-end library may be provided optionally to supplement assembly graph links.

Binning refining certainly depends on the quality of the initial binning being refined: no refining procedure could “invent” new bins. In order to reduce the variation of the results that might depend on the initial binning we used three state-of-the-art binners MetaBAT2 [10], MetaWrap [37] (which internally bins using MetaBAT2, CONCOCT and MaxBin2 and produces some sort of consensus binning) and VAMB [25] to produce three initial binnings for METAMVGL and BinSPreader. Because Binnacle is compatible with a limited number of binners, we used it with MetaBAT2 only. Unless stated otherwise, input metagenomic assembly graph was constructed using metaSPAdes 3.15.3 [26].

Resulting binnings of mock and simulated samples were analyzed with AMBER [18]. AMBER assessment of bin quality is based on annotation of metagenomic contigs using the reference genomes provided as a “gold standard binning”. Contig alignment to reference genomes was performed using metaQUAST [19]. Evaluation of real metagenomes without references were done via CheckM [27]. AMR genes were searched using RGI 5.2.1 with CARD database 3.1.4 [16]. CRISPRs were detected using MinCED 0.4.2 [1]. rRNA were annotated with Barrnap 0.9 [31].

### 2.3 Completeness, contamination and F1

In order to benchmark BinSPreader against state-of-the-art binning refining tools, namely METAMVGL, MetaCoAG, and Binnacle, we analyzed the average (mean) purity, completeness and F1-score of the binning results calculated by AMBER (at the nucleotide level) for four synthetic datasets. To complement these metrics we also took into account the number of recovered high-quality genomes with > 90% completeness and < 5% contamination as reported by AMBER. Individual F1-scores for refined bins across all datasets can be found in Supplementary Figures (1, 2, 3, 4).

On **magsim-MGE** dataset MetaBAT2, VAMB and MetaWRAP recovered very pure bins with average purity taking values from at least 97.2% for MetaBAT2 to 99.9% for VAMB and MetaWRAP (refer to Supplementary Table 1 for all AMBER metrics of this dataset). Yet these binnings had very low average completeness with maximum value of 69.2% for MetaBAT2 and minimum of 43.5% for VAMB. This poor trade-off between purity and completeness is indicated by the moderate values of the mean F1 score. Best-performing binning tool MetaBAT2 resulted in F1 score of 80.8% and recovered 12 high-quality out of 30 total genomes, the worst-performing tool was VAMB with an F1 score of only 60.6% and 8 recovered genomes.

Although refining of initial bins with METAMVGL and BinSPreader led to a minor decrease in average bin purity (no more than 3% for METAMGVL and 1% for BinSPreader across all bins), it significantly reduced the number of unbinned contigs and increased average bin completeness. Bins refined with METAMGVL and BinSPreader had average completeness ranging from 50% for VAMB and MetaWRAP to 72% for MetaBAT2. Refining MetaBAT2 bins using Binnacle did not affect bin purity compared to running MetaBAT2 alone, but reduced average completeness. MetaCoAG produced bins with average purity of 97.5%, average completeness of 47.3%, F1 score of 63.7% and 10 high-quality MAGs yielding results somewhat worse than several standalone binners.

Of all binning and refining approaches MetaBAT2 bins refined using BinSPreader with paired-end reads showed the best average F1 score of 85.0%, although metaWRAP bins refined using BinSPreader contained more high-quality MAGs (14 for MetaWRAP + BinSPreader vs 12 for MetaBAT2 + BinSPreader).

Available data of **simHC+** dataset allowed benchmarking the performance of BinSPreader against Meta-CoAG only (refer to Supplementary Table 2 for all AMBER metrics) since no original paired-end reads were available in MetaCoAG paper and therefore one cannot run METAMVGL or Binnacle using only assembly graph and provided abundance profiles. For initial binnings we used VAMB bins as well as pre-computed bins of MaxBin2, MetaBAT2. The initial bins had the average F1 scores 23.0%, 84.5%, and 91.7% for MetaBAT2, MaxBin2 and VAMB, respectively. Poor value of F1 score for MetaBAT2 binning is a result of 13.0% average bin completeness which is the lowest among all binners. Refining of MetaBAT2 with BinSPreader overall increased bin completeness to 88.4% and F1 score to 76.3% but caused a major drop in average purity of bins. VAMB showed the best balance between precision and sensitivity, although many of the contigs remained unlabeled by VAMB. Refined with BinSPreader VAMB bins showed the increase of the F1 score value to 94.1% and the number of high-quality MAGs increased from 56 to 61. MetaCoAG showed somewhat lower F1 score of 86.7% and captured only 43 high-quality genomes, therefore BinSPreader + VAMB is the best-performing pair for the **simHC+** dataset.

Binning assessment of **Zymo** mock metagenome showed 100% average purity of MetaBAT2, VAMB, and MetaWRAP bins (refer to Supplementary Table 3 for more details). Among these VAMB produced bins with the highest average completeness of 96.5% and the highest value of F1 score of 98.2%. MetaWRAP and MetaBAT2 recovered bins with poorer completeness of 78.8%, 66.2% and moderate F1 scores of 88.1% and 79.7%, respectively. Refining of MetaBAT2 bins with Binnacle decreased the value of average completeness down to 60.6%. Refining with METAMGVL led to decrease of the purity of bins down to 88.4% for MetaBAT2 and no visible changes of VAMB and MetaWRAP bins. MetaCoAG showed better trade-off between precision and sensitivity of binning yielding 85.0% F1 score but labeled fewer contigs than BinSPreader. BinSPreader significantly increased bin completeness with negligible effect on purity value that is demonstrated by F1 scores of 87.6% of refined MetaBAT2 bins, 97.5% of MetaWRAP and 99.7% of refined VAMB bins. Supplementing BinSPreader with paired-end library allowed the increase of F1 score up to 100% on VAMB bins achieving the best binning result for **Zymo** dataset.

Binning results for the **MBARC26** mock community are described in Supplementary Table 4. Initial binnings showed balanced precision and sensitivity with average F1 value of 89.4% for MetaWRAP-produced bins and 93% for VAMB and MetaBAT2. Refined bins produced by METAMVGL had lower quality than initial binning of the MetaBAT2, VAMB and MetaWRAP alone. F1 score of bins recovered with Binnacle and MetaBAT2 dropped from 93.2% down to 89.1%.

MetaCoAG showed better performance with 93.9% average purity, 92.6% average completeness and F1 score of 93.2%. F1 scores of BinSPreader refining of MetaBAT2 and VAMB bins were 94.7% and 94.5%, respectively. BinSPreader had a major impact on MetaWRAP binning quality, raising average completeness from 80.0% to 98.9% and decreasing an average purity from 99.8% to 92.3%. This binning approach showed the highest value of F1 score of 95.5% among all tested tools.

Finally, we benchmarked BinSPreader on **BMock12** mock dataset (refer to Supplementary Table 5 for all AMBER metrics). Bins of initial binning tools had high average purity ranging from 96.5% for MetaWRAP to 98.1% for VAMB and moderate average completeness taking values from 66.9% for MetaBAT2 to 79.3% for MetaWRAP. The F1 scores were in the interval from 79.4% (MetaBAT2) to 87.1% (MetaWRAP). MetaCoAG bins had lower average bin purity of 88.6% and correspondingly lower F1 score of 81.3%. Refining of bins produced with MetaBAT2, VAMB, and MetaWRAP using METAMVGL and refining of MetaBAT2 with Binnacle both led to considerable decline of all metrics as compared to the original bins. METAMVGL refining of VAMB bins resulted in 9% less average purity and 8% less average completeness compared to the initial VAMB bins. Of all refining tools, only BinSPreader effectively improved the quality of an input binning. Average F1 scores of MetaBAT2, VAMB and MetaWRAP bins refined using BinSPreader had values of 89.5%, 94.3%, and 94.6%, respectively. MetaWRAP + BinSPreader also retrieved 7 high-quality MAGs out of 11 total genomes, more than any other of the tools tested.

Summarizing the results on all datasets, graph-aware refiners METAMVGL and Binnacle either yield no noticeable effect (**magsim-MGE**) or impaired the characteristics of the original binning (**MBARC26**, **BMock12**, **Zymo**). MetaCoAG showed a decent ratio of precision to sensitivity but left large portions of contigs unbinned. Exploiting the assembly graph to the fullest extent allowed BinSPreader to augment the bins with unbinned contigs and improve their F1 score with the best trade-off between completeness and contamination. Moreover, it also increased the number of complete MAGs represented with minimal contamination.

We need to outline that the performance of any binning refining tool including BinSPreader depends on the quality of the input bins as the refiner cannot “invent” e.g. a missed bin. This pitfall is demonstrated on BinSPreader refining of the **simHC+** binning by MetaBAT2. Due to the extremely low completeness of the initial binning BinSPreader failed to accurately perform contig labeling that caused additional contamination of the bins.

In order to benchmark BinSPreader on real **IC9** and **Sharon** datasets, we used mean purity, completeness, and F1-score metrics, which were assessed using CheckM [27], as well as total number of bins. Individual F1-scores for refined bins for **IC9** and **Sharon** can be found in Supplementary Figures 10 and 11, respectively.

As reported in Supplementary tables 7 and 8, MetaWRAP showed the best average F1-score among the initial binners for both **IC9** and **Sharon** datasets (96.5% for IC9 dataset, 98.3% for Sharon). None of the graph-based refiners, namely BinSPreader, METAMVGL, and Binnacle, showed any significant improvement upon initial binnings for both real datasets, with the exceptions of BinSPreader complemented with Hi-C reads for MetaBAT2 on IC9 dataset (64.7% average F1 score for MetaBAT2 against 69.6% average F1 for BinSPreader), and Binnacle-refined MetaBAT2 binning for Sharon dataset (81.3% for Binnacle against 76.6% for MetaBAT2). DAS_TOOL refining demonstrated the best increase in average F1-score for all initial binnings. This, however, could be explained by consistent decrease in the number of bins after DAS_TOOL refining due to filtering out bins with poor CheckM metrics. Specifically, MetaBAT reported 50 bins for **IC9** dataset, while DAS_TOOL reported only 23 refined MetaBAT2 bins.

Negligible increase of CheckM purity and completeness metrics after graph-based refining for real datasets could be explained by limitations in CheckM single-copy gene-based purity and completeness estimation (they are essentially located on long contigs that are likely properly binned and no shorter contigs contribute to these metrics) and by segmentation of metagenomic assembly graphs constructed for these datasets. Indeed, for **Sharon** and **IC9** datasets the mean number of links outgoing from an assembly graph edge is 1.62 and 0.51, respectively, while for mock **Zymo** dataset the mean number of outgoing links is 2.71. Also the bins seem not to cover the whole assembly (30-60% depending on the binner).

Still, even sparse assembly graphs provide BinSPreader with sufficient information to reconstruct different functional genes more efficiently compared to initial binning as we show below.

### 2.4 Conservative genes recovery

Efficient binning of rRNA still remains one of the greatest challenges in metagenomics as rybosomal RNA gene clusters are hard to assemble due to a high number of intra- and interspecies repeats. Consequently, contigs containing rRNA genes are usually small and belong to multiple genomes. Most of the binners do not support assignment of one contig to multiple bins making it nearly impossible to recover sufficiently complete set of rRNA genes for more than one genome, even if rRNA genes were lucky to be assembled completely. We show how BinSPreader ability to propagate bin labels to small contigs and repeat regions as well as multiple bin assignment could help in rRNA recovery. Beyond that, this approach could also help in genomic islands (GI) recovery that contain regions that are important for clinical applications such as CRISPRs and antimicrobial resistance (AMR) genes.

CRISPRs (Supplementary Table 9) are not very well assembled in **MBARC26** and **magsim-MGE** datasets, as 18% and 28% of them, respectively, are missing from the assemblies. Nevertheless, BinSPreader shows the best performance recovering all repeat clusters for mock datasets regardless of refining mode. All standalone binners recover nearly equal amount of CRISPRs, but MetaCoAG manages to greatly surpass them on **MBARC26** (42 recovered CRISPRs against 33 for the best initial binner, MetaWRAP).

However, the most interesting dataset in terms of GI recovery is **magsim-MGE** as it was specifically designed to showcase this problem [14]. Refining with BinSPreader using assembly graph alone does not significantly increase the amount of recovered CRISPRs, but the usage of supplementary paired-end connectivity information gives one of the best results among all binners and BinSPreader runs particularly well (17 recovered CRISPRs out of 23 total assembled versus 13 without paired-end reads). On this dataset METAMVGL manages to recover similar number of CRISPRs as BinSPreader.

The results of AMR genes recovery (Supplementary Tables 10, 11) are pretty much consistent with CRISPRs recovery. BinSPreader and MetaCoAG still show the best performance, recovering every single assembled AMR gene on mock datasets. In contrast with CRISPRs results, running BinSPreader with paired-end information on **magsim-MGE** dataset yields the best result with MetaBAT2 as initial binner (138 recovered CRISPRs out of 145 assembled), while the number of recovered AMR genes after refining with METAMVGL was lower compared to initial MetaBAT2 binning (108 recovered genes after refining vs 115 original AMR genes).

The influence of supplementary connectivity information on the binning refining productivity can be seen on **IC9** dataset, where Hi-C data is available in addition to paired-end reads (Supplementary Table 11). BinSPreader provided with Hi-C links recovered the maximum amount of AMR genes among all binners and refiners (191 recovered AMR gene out of 300 assembled). This result could be explained by presence of Hi-C links between chromosomes and plasmids harboring AMR genes, allowing BinSPreader to propagate bin labels to plasmidic contigs more accurately.

While the amount of recovered GI and functional elements appears to be an informative benchmark for metagenomic studies, the final goal of most researches is to get as much high quality MAGs containing all these elements as possible. In order to make a high-level assessment of MAG recovery, we applied MAG reporting standards developed by the Genomic Standards Consortium [2]. MIMAG standard uses different levels of genome completeness and contamination as well as rRNA gene presence. Depending on these metrics MAGs are divided into several groups including Medium-quality draft (≥ 50% completeness, <10% contamination) and High-quality draft (>90% completeness, <5% contamination, full set of rRNA genes and at least 18 tRNA). Since rRNA recovery is primarily limited by its complete assembly, we constructed perfect binning from input assemblies that comprises MAGs with 100% purity and 100% completeness to use it as reference. We also added the second type of High-quality MAGs somewhat lowering the standard: we require a complete set of 16S or 18S rRNAs as these particular rRNA genes are of most importance for further taxonomic annotation.

Results obtained for **Zymo** and **BMock12** datasets (Supplementary Figures 12, 13) emphasize that the assembly quality plays a crucial role in rRNA recovery. Only one High-quality MAG could be obtained from **BMock12** assembly due to the fragmentation of rRNA gene contigs and only 2 High-quality MAGs (including only 16S rRNA) could be recovered from **Zymo** (Supplementary Tables 12, 14) in general. Still, BinSPreader was able to recover these MAGs from VAMB bins with the help of supplementary paired-end connectivity information. Also BinSPreader refining enriches MetaBAT2-produced bins with medium-quality MAGs (Supplementary Figure 12) for **Zymo** dataset.

On **MBARC26** and **magsim-MGE** datasets (Supplementary Figures 14, 15) we can observe a great improvement in High-quality MAG recovery after the refinement with BinSPreader in multiple binning mode. In comparison with initial bins, BinSPreader refining clearly led to saturation of MAGs with rRNA genes and other small contigs, rather than increasing a number of medium-quality MAGs. The usage of multiple binning approach increases a number of high quality MAGs almost down to assembly level.

Particularly, refining of VAMB binning of **MBARC26** dataset resulted in recovery of all 4 possible high quality MAGs. Different variations of BinSPreader modes yield 1 high quality MAG with the full set of rRNA in the worst case, which is still unattainable for the most binners, moreover all BinSPreader runs increased a number of high quality MAGs containing only 16S rRNA dramatically, especially when multiple bin assignment mode was used. Even greater improvements could be observed in refining of binning results obtained on **magsim-MGE** dataset. BinSPreader manages to recover all high quality MAGs using metaWRAP and VAMB bins without losing any medium quality MAGs. In addition, BinSPreader recovers 16S rRNA for almost for every MAG in VAMB and MetaWRAP-produced bins. Refining MetaBAT2-produced bins using paired-end connectivity information leads to recovery of five new medium quality MAGs.

On the real **IC9** metagenome, BinSPreader retrieved all 16S and 23S rRNA genes present in the assembly regardless of initial binning and genome fraction (GF) as shown in Supplementary Table 16, while the second-best refiner-binner combination, bin3C + DAS_TOOL, reconstructed only 4 23S rRNA out of 6 and 2 16S rRNA out of 3 (for rRNA genes assembled at 90% GF). Overall, BinSPreader recovered 71 rRNA genes out of 73 (against 36 for the next best refiner, MetaCoAG). On the **Sharon** dataset BinSPreader supplemented with paired-end reads retrieved 20 out of 29 of all rRNA genes assembled with at least 50% GF, while second best refiner, MetaCoAG, recovered only 6 rRNA genes (see Supplementary Table 17).

### 2.5 Binning refining supplemented with paired-end and Hi-C linkage

To assess the effectiveness of paired-end reads information for binning refining, we used paired-end read libraries available for **Zymo**, **MBARC**, **Bmock12**, and **magsim-MGE** datasets. We compared MetaBAT2, VAMB, and MetaWRAP bins refined with BinSPreader supplemented with paired-end reads (*BSP-PE mode*) and bins refined with BinSPreader provided with assembly graph only (*BSP mode*). We also assessed Binnacle and METAMVGL refiners which utilize paired-end reads as well. We evaluated binning results using AMBER [18] and reported F1-score of the initial and refined bins.

For **magsim-MGE** dataset, Supplementary Table 1 shows that BSP-PE results in higher F1-scores than BSP for all three initial binners. For **Zymo** dataset, Supplementary Table 3 shows that BSP-PE resulted in higher F1-score per sample than BSP for VAMB and MetaBAT2 binnigs (87.6% for BSP-PE versus 86.7% for BSP for MetaBAT2, 100% for BSP-PE versus 99.8% for BSP for VAMB), and the same F1-scores for MetaWRAP binning. For **BMock12** dataset, BSP resulted in higher F1-score for MetaBAT2 and MetaWRAP datasets than BSP-PE, but BSP-PE for VAMB binning showed the highest F1-score across all binners and refiners (94.6% for BSP-PE versus second highest 94.2% for BSP), as shown in Supplementary Table 5. For **MBARC** dataset, BSP-PE resulted in lower F1-scores than BSP for all three initial binners (Supplementary Table 4). The possible reason for this is contamination in paired-end library for **MBARC**, since applying METAMVGL and Binnacle to all three initial binnings resulted in lower F1-score (Supplementary Table 4). For all samples and all initial binners, BSP-PE resulted in higher F1-scores than METAMVGL and Binnacle. F1-scores for separate bins are reported in Supplementary Figures 1, 2, 3, 4.

The potential of Hi-C technology as a means to cluster metagenomic contigs into bins has been demonstrated on both synthetic and real microbial communities [5, 6, 8]. We followed two approaches to analyze possible integration of Hi-C technology and binning refining methods for MAG recovery.

First, we obtained initial binning for **Zymo** Hi-C library using dedicated Hi-C bin3C [5] binning tool and refined bin3C binning using BinSPreader (in both BSP and BSP-PE modes). As shown in Supplementary table 6, F1-scores reported by AMBER were higher for bin3C bins refined by BinSPreader (0.927 for BSP and BSP-PE against 0.865 for unrefined bin3C bins).

Second, we used **Zymo** Hi-C links as an additional source of information for BinSPreader (*BSP-HiC mode*) and benchmarked the results against BSP-PE and BSP modes for MetaBAT2, MetaWRAP, and VAMB bins. For MetaBAT2 binning, BSP-PE showed highest F1-score (0.911), followed by BSP-HiC (0.903), and BSP (0.896). For MetaWRAP and VAMB binnings, BSP, BSP-PE, and BSP-HiC resulted in similar F1-scores.

While BSP-HiC did not show any improvement upon BSP-PE in terms of standard contamination and completeness metrics for **Zymo** dataset, AMR gene detection results for plasmid-rich **IC9** dataset described above (see Section 2.4) show that BSP-HiC can be used to reconstruct additional functional elements located on the unbinned contigs that were not connected to the main genome on the assembly graph.

### 2.6 MAG distance estimation using prob Jaccard index

Sometimes binners produce very pure but incomplete bins (results of section 2.3 shows that usually this is the case of MetaBAT2 and MetaWRAP bins). After refining such bins tend to overlap on an assembly graph and therefore the size of such overlap could potentially be used to decide whether one need to merge certain bins. Also, overlapped labeling of the edges of assembly graph could measure possible contamination or otherwise shared genome content.

Supplementary Figure 16 shows the hierarchical clustering of bin distance information calculated from **Zymo** MetaBAT2 bins. One could easily see the bins of different genomes clustered together as well as significant (and expected) overlap of *E. coli* and *S. enterica* bins. Supplementary Figure 17 shows the hierarchical clustering of bin distance information calculated from **BMock12** MetaBAT2 bins. Again one could see several bins of the same species located together on the graph as well as significant bin overlap between two *Micromonospora* strains as well as contamination of *Marinobacter* bins.

## 3 Methods

### 3.1 From scaffold binning to edge binning

Most binners output its results in a form of *scaffold binning,* i.e., a map *B* from a set of scaffolds *P* to a set of bins *C*. This representation is not entirely accurate, since long scaffolds in a metagenomic assembly may contain repetitive regions, which can belong to multiple species in a sample, and therefore in multiple bins. To alleviate this, BinSPreader transforms the initial scaffold binning to the *edge binning* using assembly graph. Let *G* be an assembly graph in [GFA format](https://github.com/GFA-spec/GFA-spec) consisting of a set of edges *E*(*G*), links *L*(*G*) between them, and scaffolds *P*(*G*) with their corresponding paths in the assembly graph. Given edge *e_i_* ∈ *E*(*G*), let *P*(*e_i_*) ⊂ *P*(*G*) be the set of scaffolds that contain *e_i_*, and *C*(*e_i_*) ⊂ *C* be the set of bin labels of *P*(*e_i_*). For assembly graph G and scaffold binning *B*, BinSPreader transforms scaffold binning *B* to edge binning matrix *Y*, where

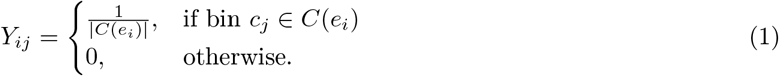

Here each row *Y_i_* represents a *soft binning* of edge *e_i_*, which can be interpreted as the containment probability distribution over the set of bins. Edge binning represents a more fine-grained representation of initial binning than scaffold binning, as repetitive edges may contain multiple bins if they are traversed by several paths (Figure 1).

**Figure 1:**
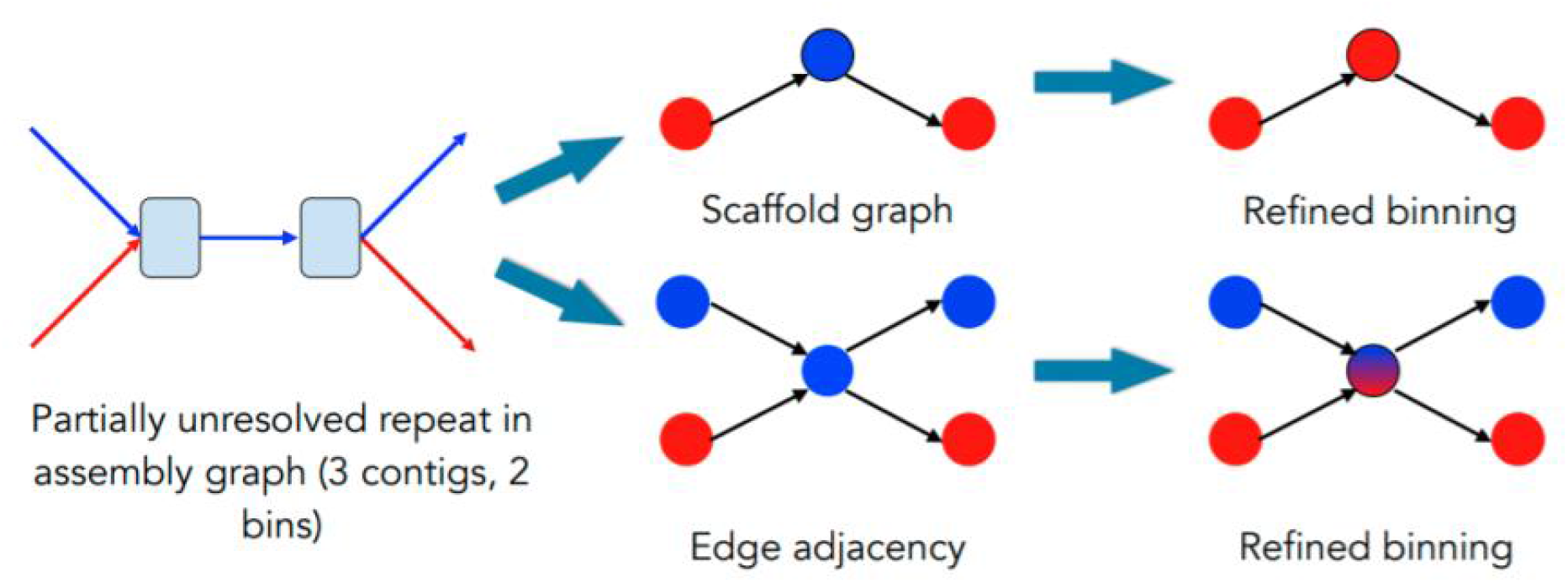
Edge adjacency graph and scaffold graph. (Left) A partially unresolved intergenomic repeat in a metagenomic assembly graph with initial binning. Three blue edges are assembled into a single contig, while two red edges belong to different contigs that were assigned into a single bin. (Top) Scaffold graph representation of the assembly graph, with vertices representing contigs and colors representing bins. Correction procedure might erroneously reassign blue vertex to a red bin based on graph connectivity. (Bottom) Edge adjacency graph representation of the assembly graph, with vertices representing assembly graph edges, and edges representing shared vertices. Repetitive edge can be correctly assigned both to blue and red bins

### 3.2 Link graph

While edges of the assembly graph *G* are used to store the initial binning and the end results, vertices of the assembly graph provide minimal required connectivity information for BinSPreader. Connectivity information is stored in a form of a weighted *link graph H*, where *V*(*H*) = *E*(*G*), *E*(*H*) = *V*(*G*) and the edge weight *L_ij_* represents the weight of a link between assembly graph edges *e_i_* and *e_j_*. The higher *L_ij_* is, the more likely is that *e_i_* and *e_j_* belong to the same bin. Initially BinSPreader uses adjacency matrix of an assembly graph *G*for weights with *L_ij_* = 1 if the edges *e_i_* and *e_j_* are adjacent in *G*and zero otherwise.

Besides the adjacency weights, BinSPreader also by default considers the set of *scaffold links*: if two edges are joined in a scaffold, but not adjacent in the graph we add the link in *H* (add edge and set *L_ij_* = 1) between them. Usually such scaffold joins are made by an assembler to jump over coverage gaps or long unresolved repeats. In both cases adding these links increases the contiguity of link graph and could help the binning propagation across assembly gaps.

In addition to the assembly graph itself BinSPreader is able to construct links from paired-end and Hi-C [13] libraries which can be provided optionally. Reads from a paired-end libraries and Hi-C libraries are aligned using k-mer alignment similar to [3]. First, we index unique k-mers in the assembly graph. Then we align a Hi-C read pair iff it contains two or more non-overlapping k-mers. We use *k* = 31 by default as most 31-mers in the metagenomic assembly graph are unique, but that value can be adjusted depending on the size of the sample. We then increase the link weight *L_ij_* by the logarithm of the total number of read-pairs aligned to *e_i_* and *e_j_* from all input libraries.

### 3.3 Binning refinement

Informally speaking, we say that an edge binning is *smooth* if soft bins associated with a pair of edges joined by a link with high weight are similar. As such, binning refining problem can be defined as finding smooth edge binning *F* which is close in some sense to the initial edge binning *Y*. Given link graph *H*, we use a quadratic form of normalized Laplacian of *H* as a standard spectral graph theory measure of smoothness [4, 23, 24]. Let *D* be a degree matrix of *H*, and *L* be an adjacency matrix of *H*. Then we define edge binning smoothness as

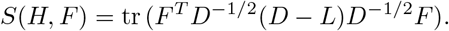

We define binning refinement problem as

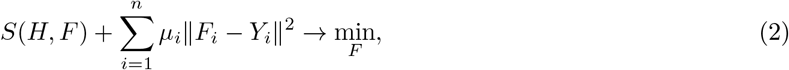

where the second term penalizes the distance between resulting binning *F* and original binning *Y* according to regularization parameters defined separately for every edge.

We use iterative algorithm for optimizing cost function (2), which is similar to one from [22]. Let 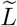 be the normalized weight matrix *D*^-1/2^(*D* – *L*)*D*^-1/2^, where *D* is a degree matrix of *H*. Then let 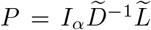, where 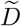 is a diagonal of 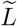, *I* is an identity matrix of size |*V*(*H*)| × |*V*(*H*)|, and *I_α_* is a diagonal matrix being *I_ii_* = 1/*μ_i_*. Initially, we set *F*(0) = *Y*. At each iteration, for every assembly edge *e_i_* the soft labels from neighboring links (*e_i_, e_j_*) with weight *H_ij_* are added to the soft label of *e_i_* with coefficient *H_ij_*. At iteration *k* + 1 we set

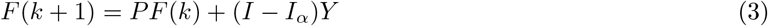

As shown in [22], the obtained sequence *F*(*k*) will eventually converge to solution 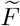, which is produced as the resulting edge binning.

We need to explicitly note that while all the matrices involved are quite large, they are extremely sparse and there is no need to store and calculate them explicitly. The soft binning for each edge at iteration k (the rows of F(k)) depends only on soft binnings of adjacent edges (which in ordinary de Bruijn graph case is not more than 8) as well as normalized link weights. This enables computational and memory efficient way to perform the iterations.

### 3.4 Choosing regularization parameters

The choice of per-edge regularization parameters *α_i_* = 1/*μ_i_* is different for different working modes of BinSPreader. Firstly, we always set *α_i_* = 1 for all repetitive edges (i.e. the edge that belongs to multiple scaffolds). As it could be easily seen from (3), the original binning for such edges will be ignored and soft binning for such edge is determined entirely via binning propagation. However, the binning from binned repetitive edges will be propagated down to their neighbors. This ensures proper and fair binning in case of e.g. partially unresolved repeats (see Figure 1 as an example).

Setting *α_i_* = 0 for edge *e_i_* would force use of original binning. This is done for all non-repetitive binned edges in *propagation mode* of BinSPreader. In such case the original binning is essentially preserved and only propagated further on to unbinned edges.

Setting 0 < *α_i_* < 1 for edge *e_i_* allows one to balance between preserving of the initial binning and propagating the binning from adjacent edges. In *correction mode* of BinSPreader *α_i_* is set to 0.6 by default for all binned edges longer than 1000 bp, for shorter edges the value of *α_i_* is gradually increasing up to *α_i_* = 1 for edges of length 1. The motivation for this is as follows: while short edges might be unique and belong only to the single scaffold, they likely repetitive and belong to unresolved repeats. The shorter the edge is, the higher its likelihood of being repetitive and we equally treat all edges longer than 1000 bp. Certainly, the latter still might be repetitive and this is what the default value of *α_i_* = 0.6 tries to accommodate.

### 3.5 Sparse binning & propagation

Binnings of real metagenomic datasets are typically sparse, since large datasets contain strains with high enough coverage to contribute to metagenomic assembly, but not high enough to be binned using the abundance and nucleotide profiles.

BinSPreader uses a special working mode of the binning refining algorithm for *sparse* binnings, where the total length of initially binned contigs is significantly lower than the total assembly length. Below we show why the standard mode of BinSPreader produces highly contaminated bins when refining sparse binnings and describe the *sparse mode* of BinSPreader designed to alleviate that problem.

Given assembly graph *G* with the set of regularization parameters *α_i_*, and initial edge binning *Y*, we say that edge *e_i_* is *refinable,* if *α_i_* ≠ 0. If initially unlabeled edge e is connected to initially labeled edge by a path of refinable edges, it eventually will be labeled after applying binning refinement algorithm to graph *G* and binning *Y*. Therefore, in the standard correction mode of BinSPreader with *α_i_* > 0 every unlabeled edge residing in the same connected component with labeled edges will become labeled after the refining. As such, refining of initially sparse (incomplete) binnings that cover only small part of *G* with *n* bins via the standard correction mode of BinSPreader will result in assigning of the majority of contigs in the refined binning to one or several of these same n initial bins potentially inflating and contaminating them.

To reduce the number of refinable edges while still allowing binning propagation, we adjust regularization parameters *α_i_* for initially unlabeled edges with *distance coefficients β_i_*, reflecting assembly graph distance to the closest initially labeled edge. Given assembly graph *G* and initial binning *Y*, let Dist(*e, Y*) be the length of shortest path in assembly graph *G* from edge *e* to the closest edge which is labeled in *Y*. We say that edge *e* is *distant,* if Dist(*e, Y*) > *D*, where *D* is distance threshold with default value 10000. To ensure that distance coefficients *β_i_* change smoothly from 1 for labeled edges to 0 for distant edges we utilize the same binning refining algorithm.

We introduce two bins, one for all labeled edges in *G* and another one for all distant edges. Then we run the binning refining algorithm as in standard correction mode of BinSPreader and set *β_i_* to the obtained weight of the first (“labelled”) bin. This makes the values of *β_i_* to gradually decrease from being 1 in case of initially binned edge *e_i_* down to to 0 when moving out of binning edges on the graph.

For sparse propagation the regularization parameters are then set as 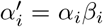, where *α_i_* are regularization parameter values for the standard correction mode of BinSPreader. This allows us to keep the initial binning intact for the edges located “far away” from the binned ones.

In addition to adjusted regularization parameters, sparse mode of BinSPreader also adds a dedicated bin for initially unbinned edges. However, while we allow the binning to propagated from binned edges down to unbinned ones we need to prevent propagation of this special “unbinned” label. In order to do so we modify the iteration procedure in sparse mode adjusting the weight matrix *P*.

### 3.6 Binning strategies: from edges back to scaffolds

After inferring refined edge binning 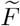, BinSPreader uses it to produce the scaffold binning *F*′. BinSPreader can output results either in single assignment or multiple assignment mode, and utilizes either *majority length* or *maximum likelihood* strategy (default).

Given a scaffold s containing edges *e*_1_,...,*e_m_*, and bin *c_j_* the binning strategy defines a score function *Score*(*s, c_j_*). For majority length strategy we define *c*(*e_i_*) = arg max_*j*_ 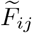 and use 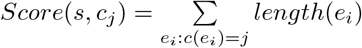.

For maximum likelihood strategy 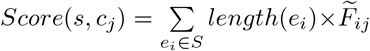. In single assignment mode BinSPreader outputs a single bin label 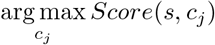 for every scaffold s. In a multiple assignment mode, BinSPreader outputs a set of labels {*c_j_*} with maximal *Score’s,* which cumulatively explain at least 95% of the total *Score.* Note that raw *Score*(*s, c_j_*) values are reported by BinSPreader as well, so one could use them for their own binning assignment procedures.

### 3.7 Measuring MAG distance using prob Jaccard Index

The typical measure to estimate the overlap of two sets is Jaccard index [9]. However, in case of BinSPreader the sets (bins) are fuzzy as the result of binning refining is a set of weights that represent the bin labeling probability distribution. In order to estimate possible overlap of bins on the assembly graph from the soft binning we consider each bin as a probability distribution on graph edges and calculate the prob-Jaccard index *J_p_* from [20] among all pairs of bins. *J_p_* has several nice features including scale invariance, it is not lower than ordinary Jaccard index valus for discrete uniform distributions (ordinary sets) and 1 – *J_p_* is a proper metric on probability distributions, meaning that *J_p_* could be used as a similarity index in e.g. hierarchical clustering and there will be no such effects like tree inversions.

### 3.8 Read extraction and MAG reassembly

In addition to providing multiple scaffold binning, accurate multiple edge binning provides an opportunity to improve upon existing metagenomic assembly using read extraction from paired-end library provided to BinSPreader. For read extraction we utilize an approach adopted from [37] from contigs down to edges. Let 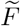 be a refined multiple edge binning and *E_j_*(*F*) be a set of assembly graph edges *e_i_* that contain bin *c_j_* with weight 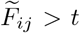, where *t* is a reassembly weight threshold with default value 0.1. We then align a set of reads from paired-end library to edges *E_j_*(*F*) separately for every bin *c_j_* obtaining set of read-pairs *R_j_*, which includes all read-pairs where at least one read aligned to *E_j_*(*F*). Such set of reads could be further reassembled or analyzed as necessary.

## 4 Discussion

Although metagenome-assembled genome binning methods based on TNF distance, coverage profiles, and single-copy marker genes are useful for untangling complex bacterial communities as a whole, they face challenges with reconstruction of functional elements located in conservative genomic regions, such as rRNAs, CRISPRs and AMR genes. This is unfortunate, given the phylogenetic and clinical relevance of these functional elements. Conservative genomic regions are usually associated with short repetitive edges of metagenomic assembly graph. Therefore, there is a clear need for metagenomic binners or refiners that enrich MAGs with short and possible repetitive contigs.

BinSPreader is a binning refining tool that effectively utilizes assembly graph connectivity information and predicts contigs belonging to several MAGs. We show that existing binning refining tools, which utilize scaffold graphs instead of assembly graphs, are less effective than BinSPreader in terms of functional element recovery (Supplementary tables 9 – 11) and in terms of rRNA genes recovery for artificial (Supplementary tables 12 – 15) and real (Supplementary tables 16, 17) metagenomes. While BinSPreader does not show significant increase in 16S/18S rRNA genes reconstruction compared to initial binning for **BMock12** and **Zymo** datasets, we show that for these datasets ability for rRNA recovery is limited mostly by assembly quality (Supplementary tables 12, 14). Experimental results on synthetic and simulated datasets show that BinSPreader also outperforms existing refiners in terms of standard contamination and completeness metrics (Supplementary Figures 1 – 4).

In addition to MAG recovery, BinSPreader provides two additional features. First, the read splitting feature, that takes into account possible overlap between MAGs and thus enables fuller MAG reconstruction after reassembly. We also introduced a bin distance measure, that provides an overlap based estimation of evolutionary distance between MAGs, thus potentially providing a novel source of information for taxonomic classification as well as detecting possible bin contamination.

## 5 Acknowledgments

The research was carried out in part by computational resources provided by the Resource Center “Computer Center of SPbU”. The authors are grateful to Saint Petersburg State University for the overall support of this work. IT and AK were supported by the Russian Science Foundation (grant 19-14-00172).

## 6 Data availability

**Zymo** and **magsim-MGE** datasets are available at https://github.com/LomanLab/mockcommunity and https://osf.io/x2y8f/, respectively. The Illumina short-reads of **MBARC**, **BMock12**, **IC9**, and **Sharon** datasets are available in NCBI Sequence Read Archive (SRA); the accession numbers are SRX1836716, SRX4901583, SRX10650162 and SRX144807, respectively. Metagenomic assemblies, references, and coverage profiles for **simHC+** are available at https://figshare.com/projects/MetaCoAG/121014. Hi-C data for **IC9** dataset is available in SRA by accession number SRX10650163. All assembly graphs, produced scaffolds, abundance profiles and binning results are available from https://figshare.com/projects/BinSPreader/132425

## Supplementary Figures

**Supplementary Figure 1:**
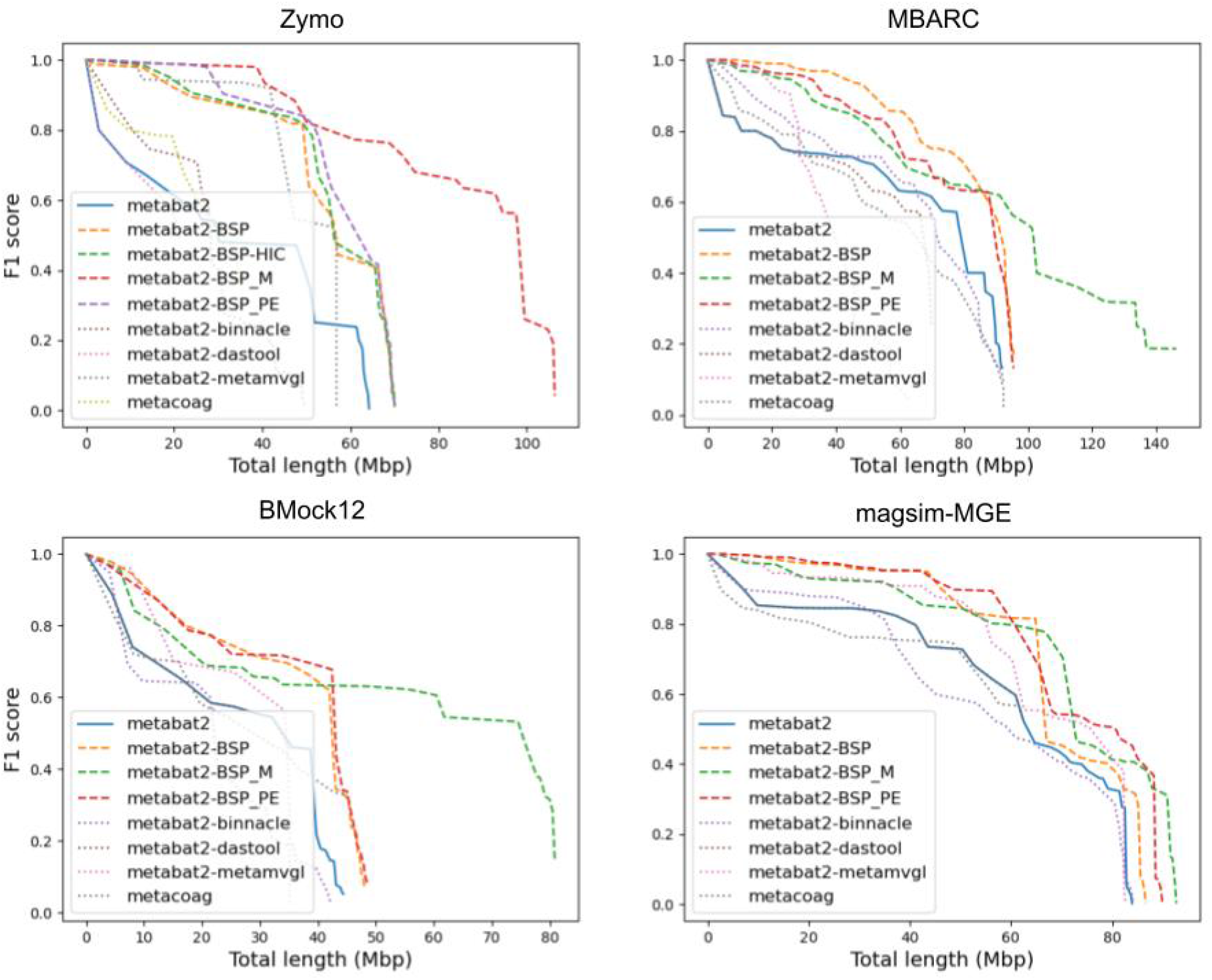
F1 score and total length of MAGs for Zymo (top left), MBARC (top right), BMock12 (bottom left), and magsim-MGE (bottom right) datasets, arranged by descending order of F1 (seq) score reported by AMBER. Initial binning was produced using metaBAT2 (solid line), refined binnings were produced using DAS_TOOL, Binnacle, and METAMVGL (dotted lines), and three different modes of BinSPreader (dashed lines).

**Supplementary Figure 2:**
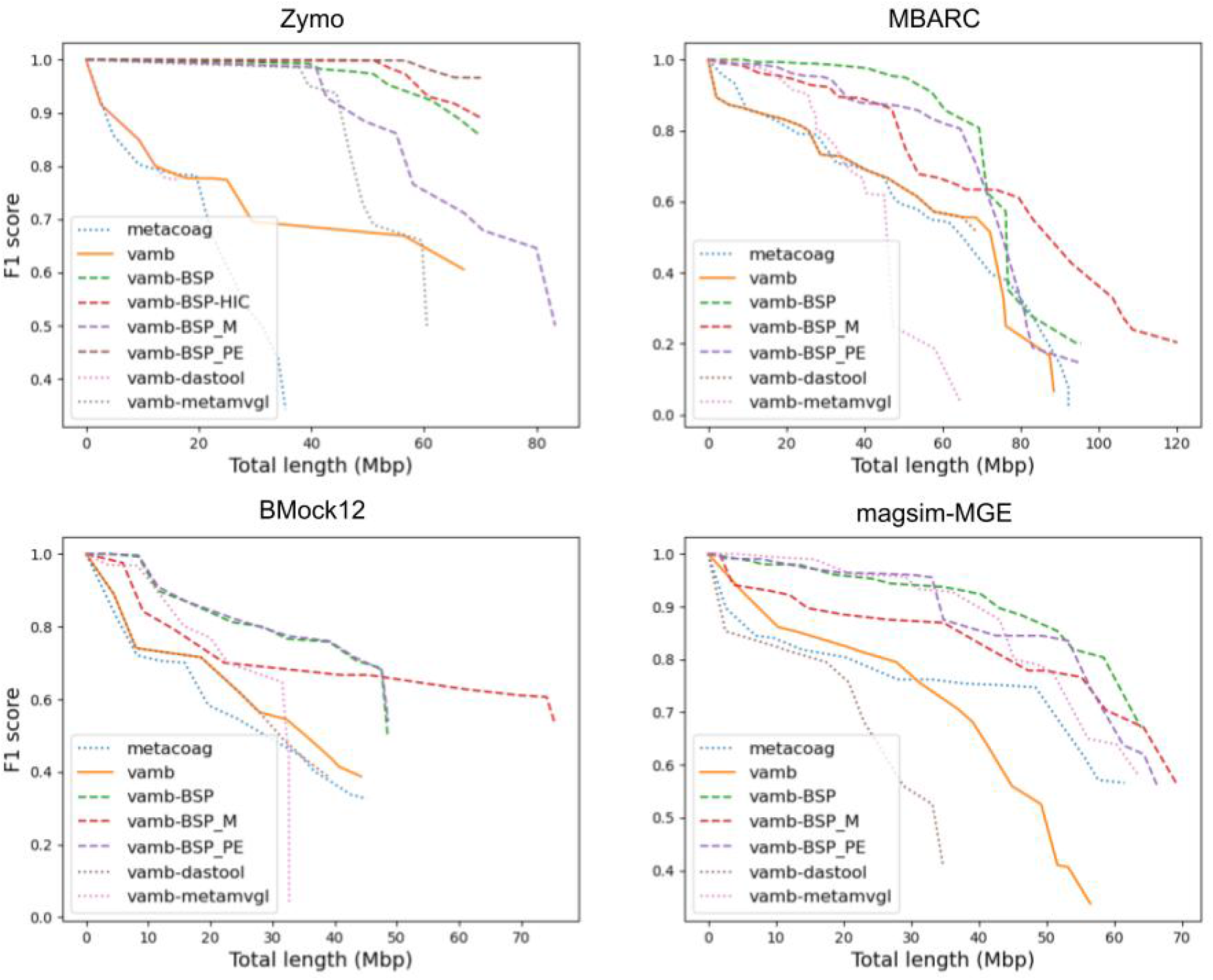
F1 score and total length of MAGs for Zymo (top left), MBARC (top right), BMock12 (bottom left), and magsim-MGE (bottom right) datasets, arranged by descending order of F1 (seq) score reported by AMBER. Initial binning was produced using VAMB (solid line), refined binnings were produced using DAS_TOOL and METAMVGL (dotted lines), and three different modes of BinSPreader (dashed lines).

**Supplementary Figure 3:**
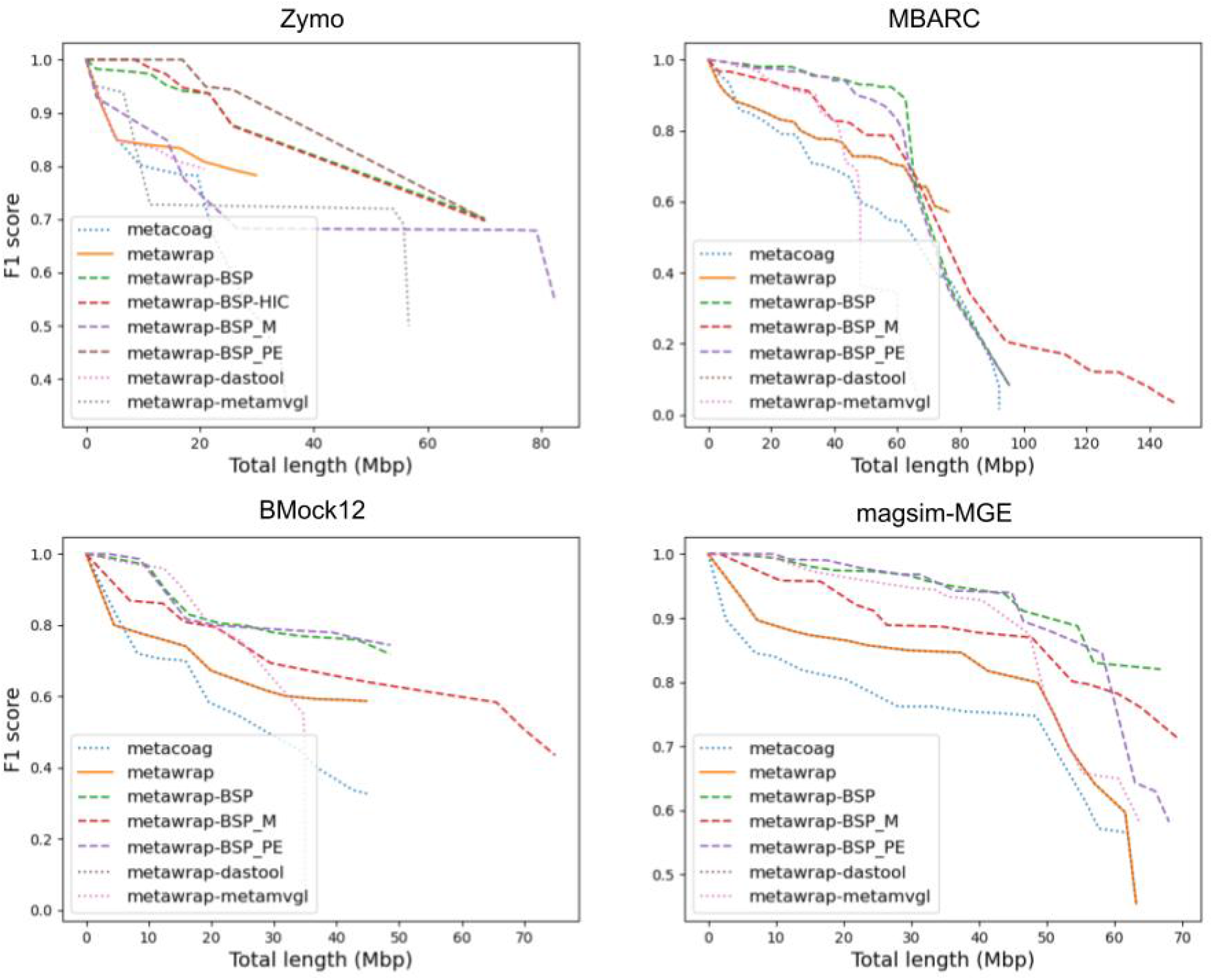
F1 score and total length of MAGs for Zymo (top left), MBARC (top right), BMock12 (bottom left), and magsim-MGE (bottom right) datasets, arranged by descending order of F1 (seq) score reported by AMBER. Initial binning was produced using MetaWRAP (solid line), refined binnings were produced using DAS_TOOL and METAMVGL (dotted lines), and three different modes of BinSPreader (dashed lines).

**Supplementary Figure 4:**
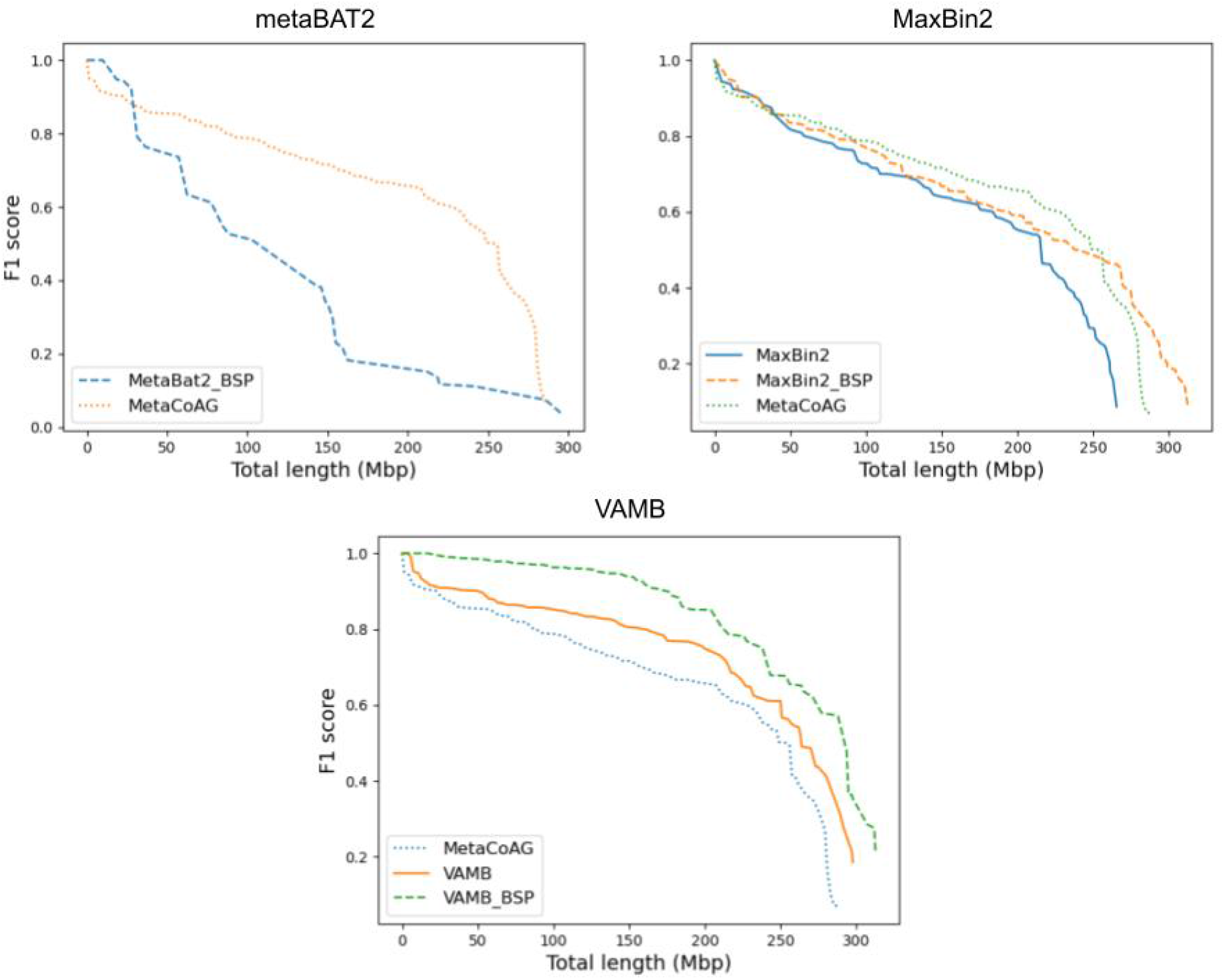
F1 score and total length of MAGs for simHC+ dataset, arranged by descending order of F1 (seq) score reported by AMBER. Initial binning was produced using MetaWRAP (top left), MaxBin2 (top right), and VAMB (bottom). Refined binnings were produced using BinSPreader (dashed line) and MetaCoAG (dotted line).

**Supplementary Figure 5:**
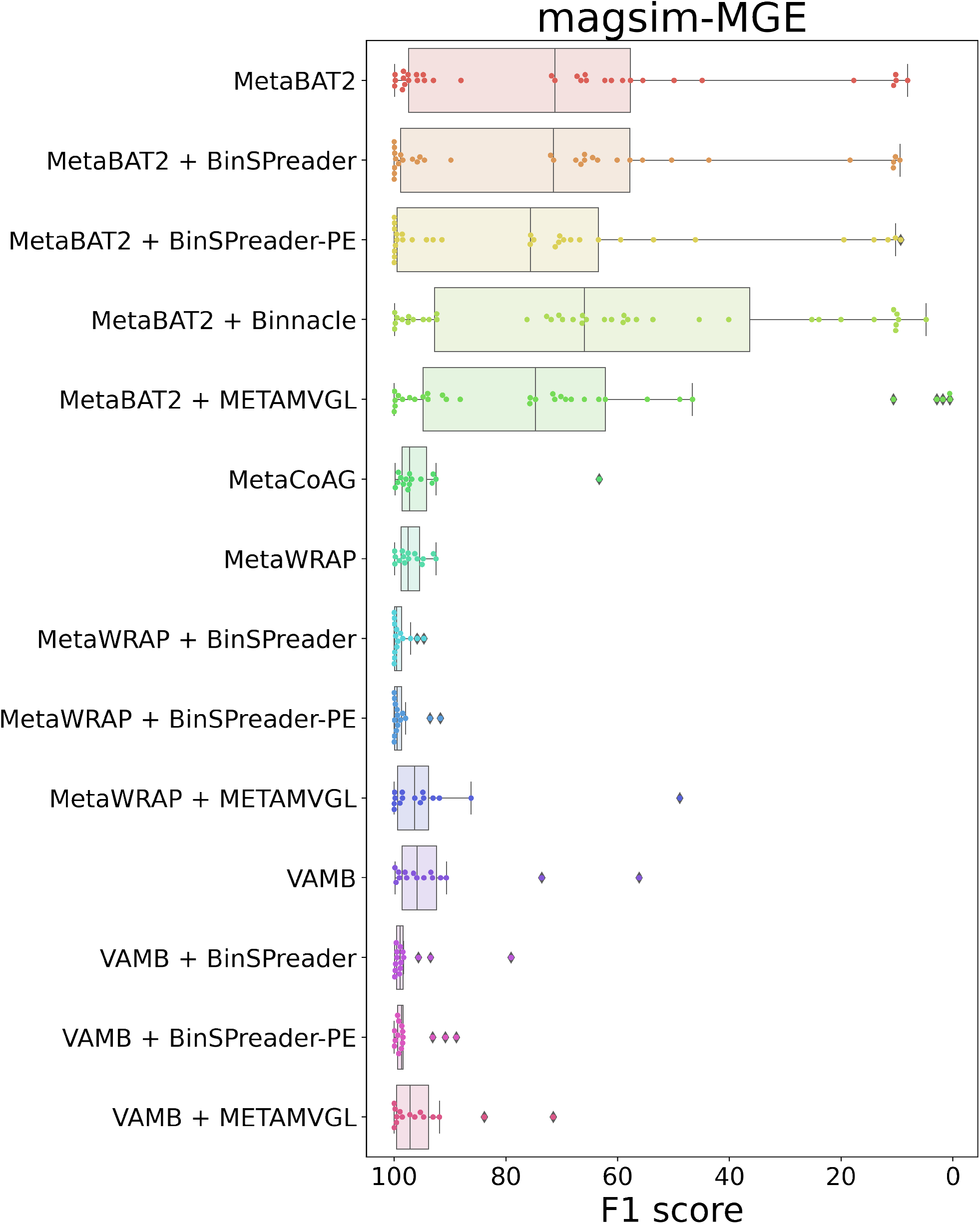
F1 score distributions of **magsim-MGE** bins.

**Supplementary Figure 6:**
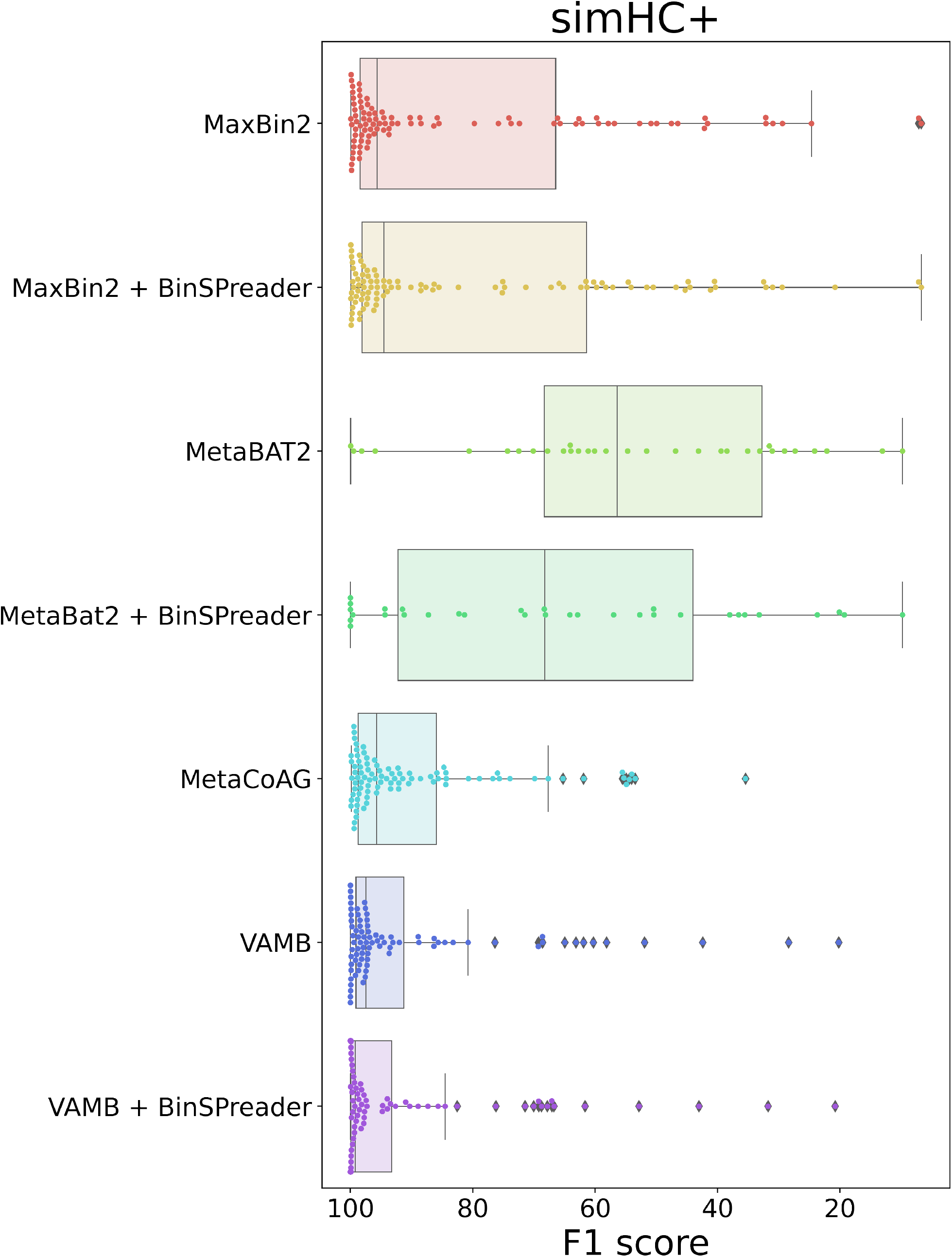
F1 score distributions of **simHC+** bins.

**Supplementary Figure 7:**
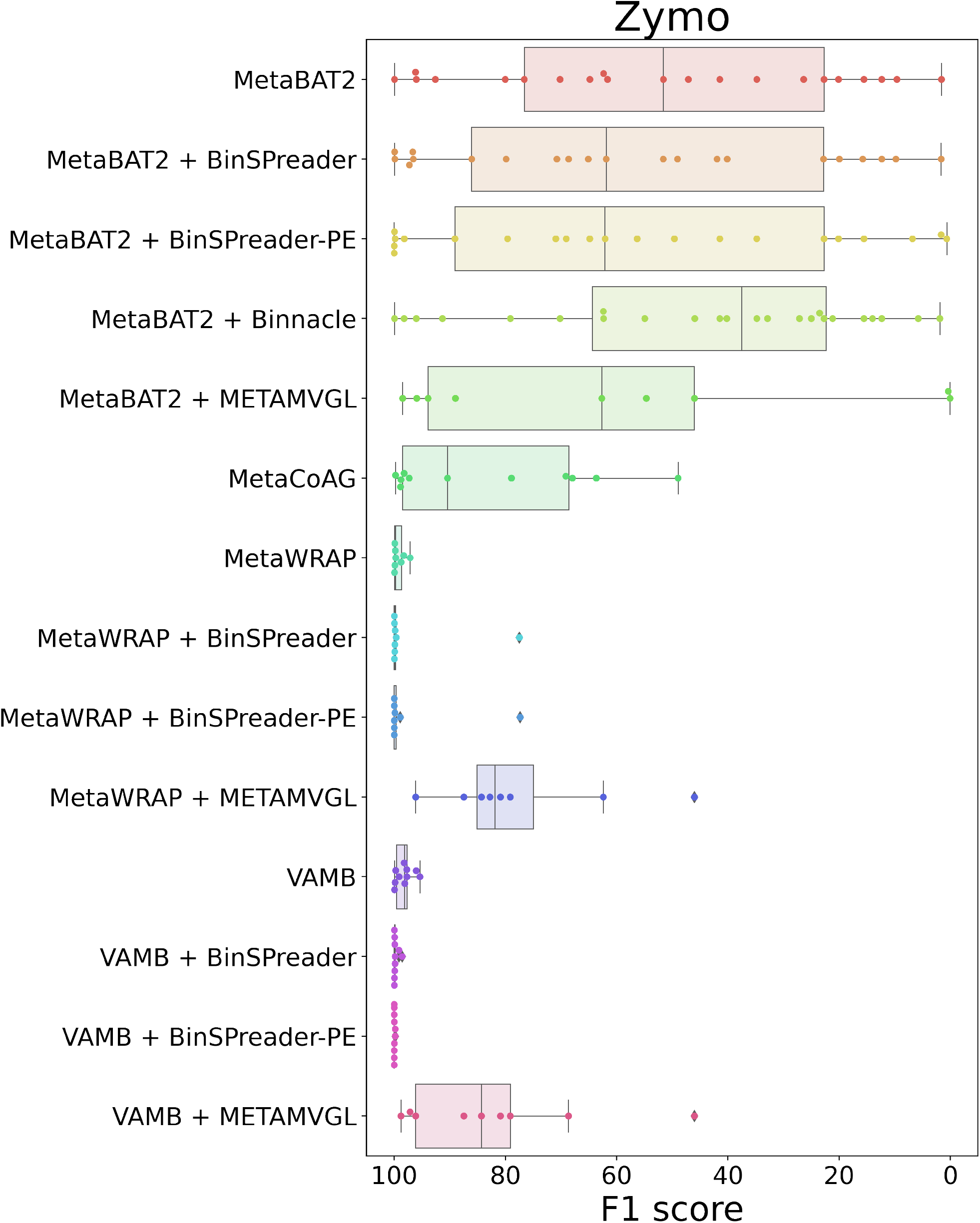
F1 score distributions of **Zymo** dataset.

**Supplementary Figure 8:**
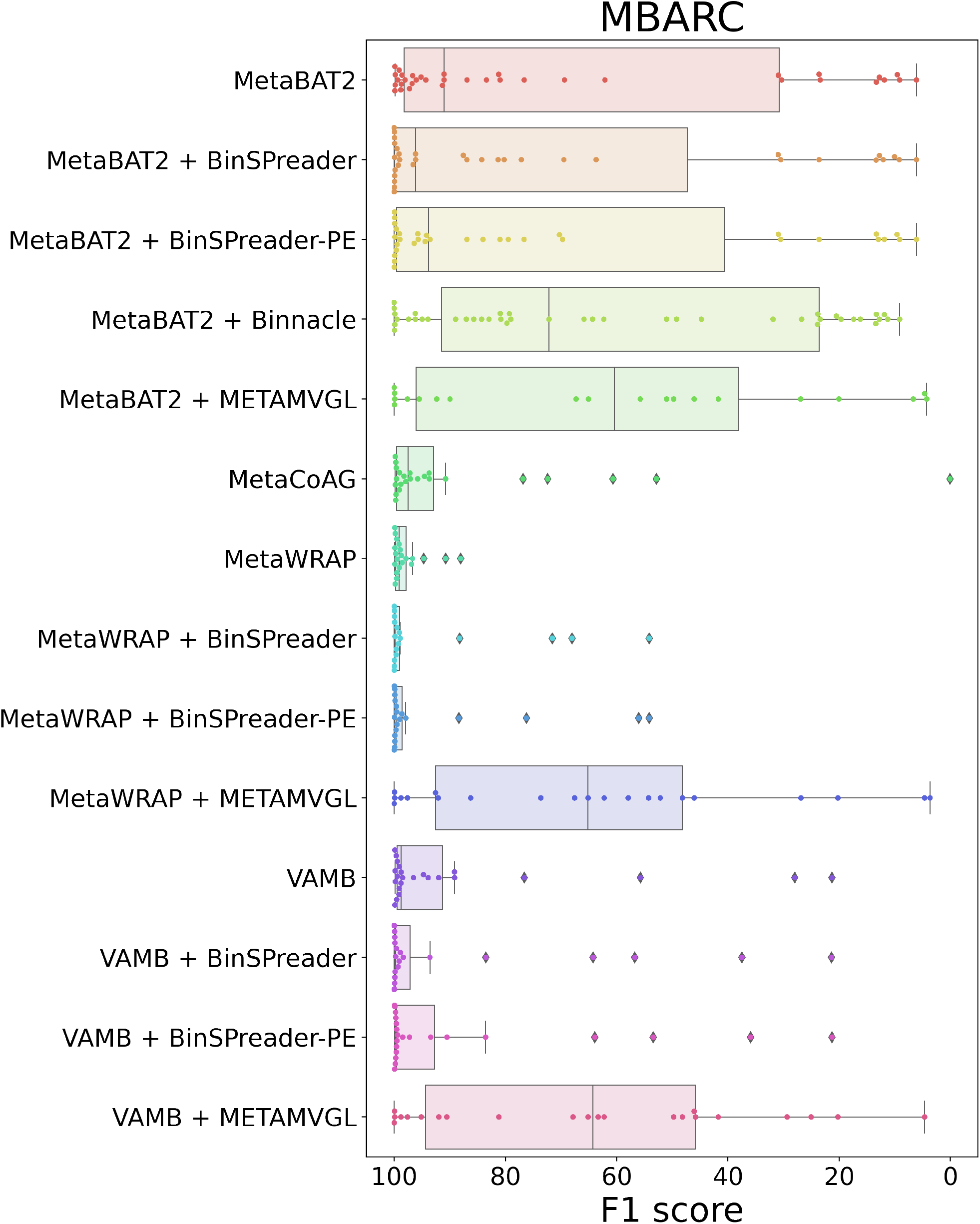
F1 score distributions of **MBARC26** bins.

**Supplementary Figure 9:**
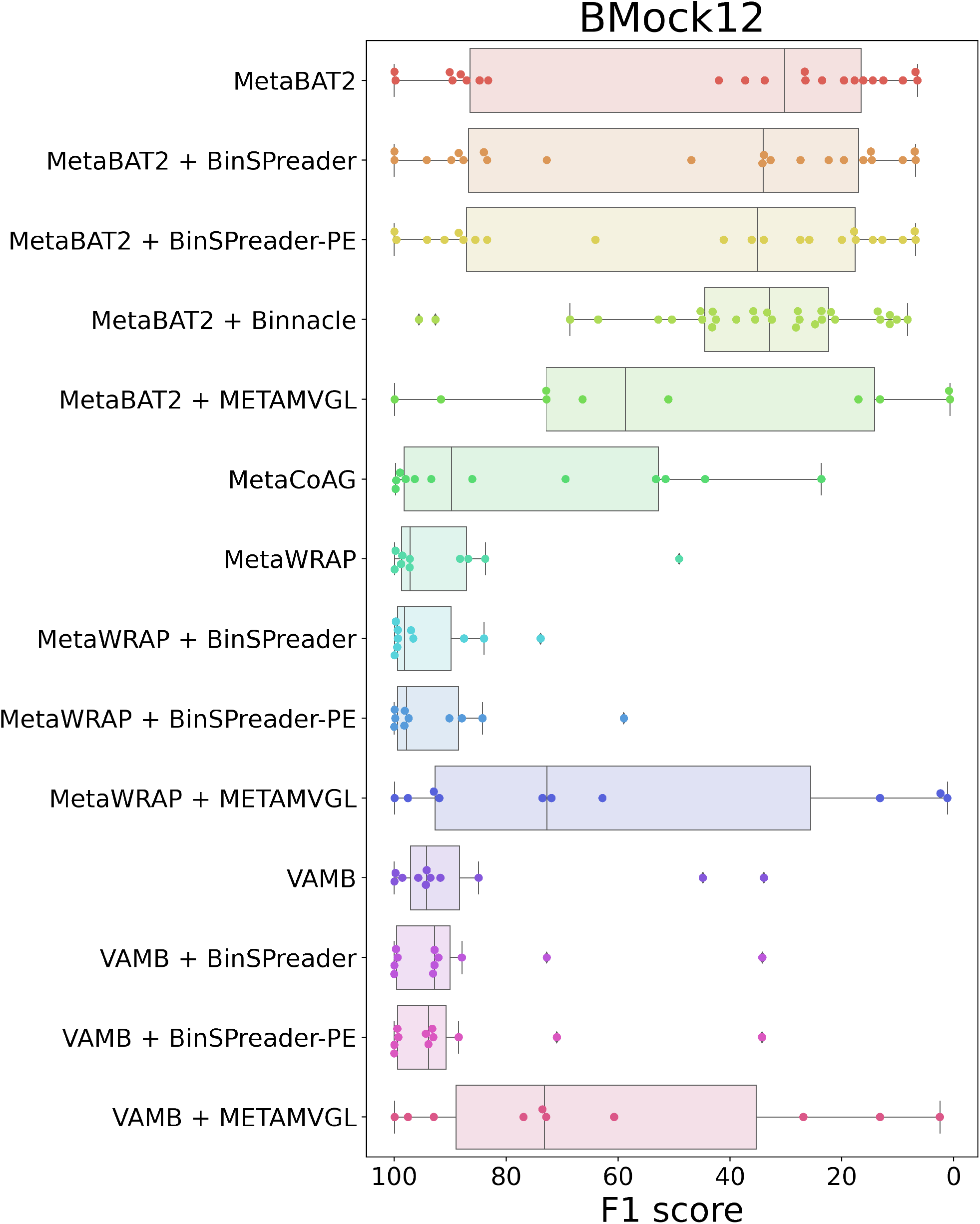
F1 score distributions of **BMock12** bins.

**Supplementary Figure 10:**
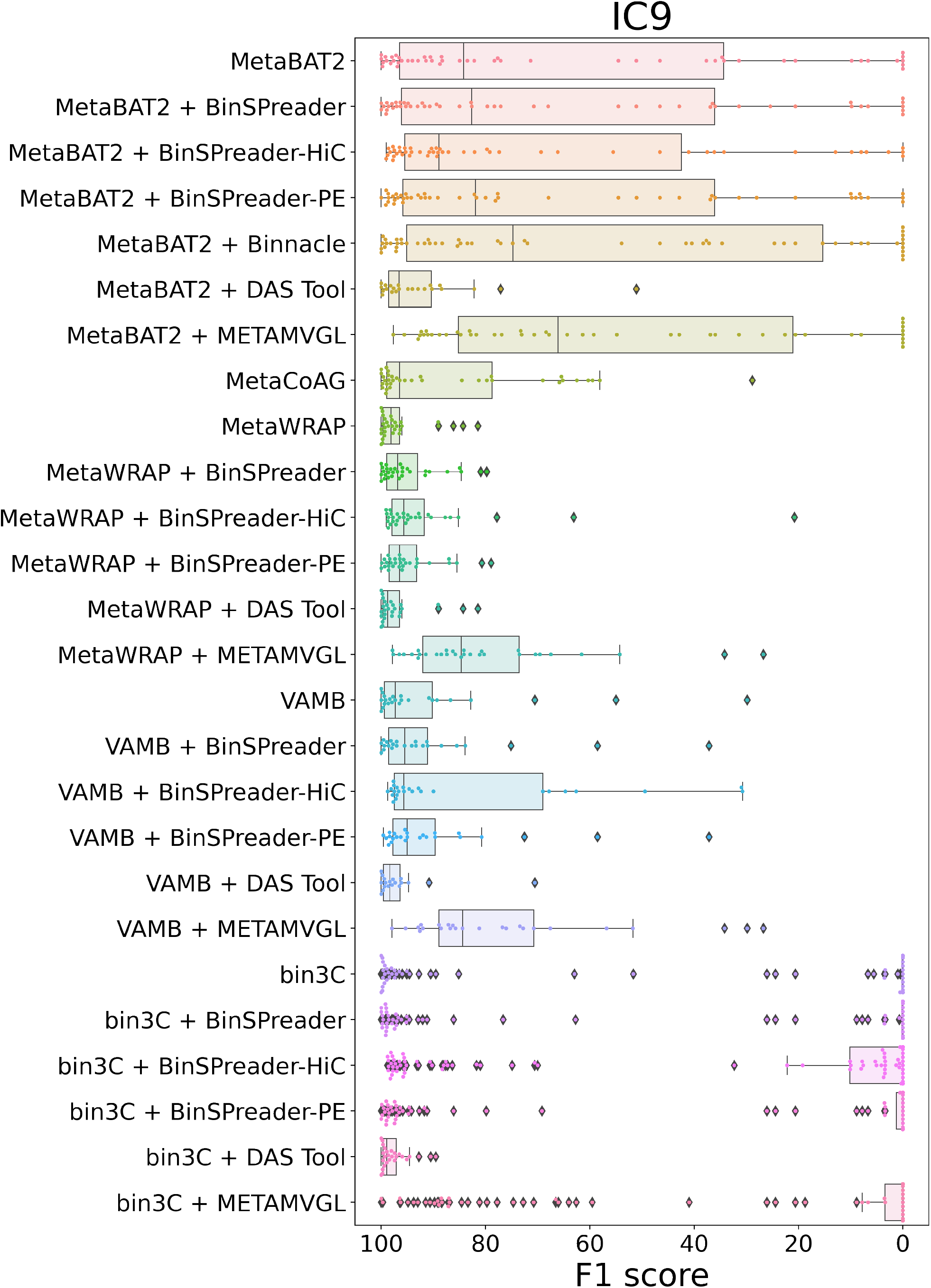
F1 score distributions of **IC9** bins.

**Supplementary Figure 11:**
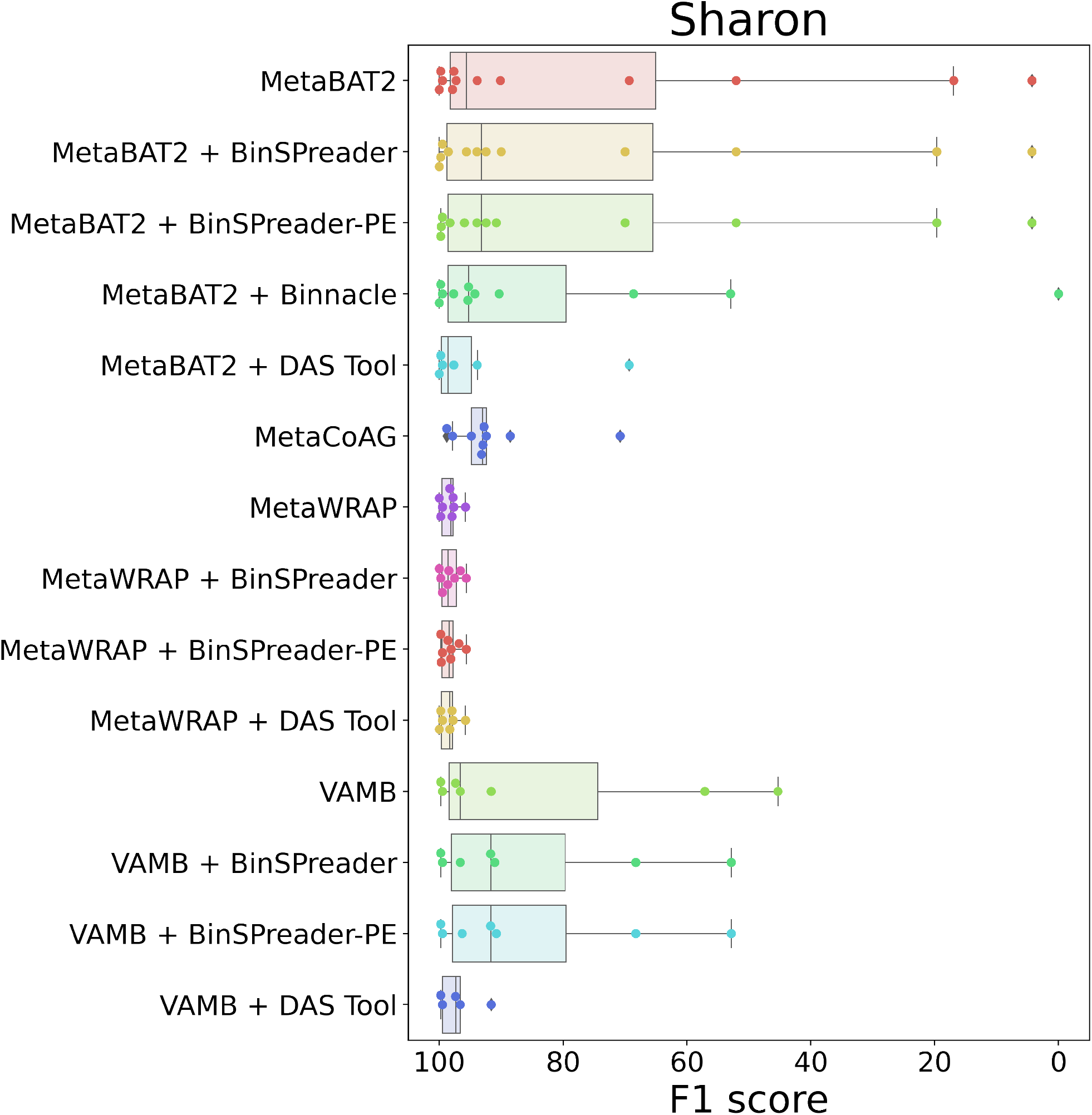
F1 score distributions of **Sharon** bins.

**Supplementary Figure 12:**
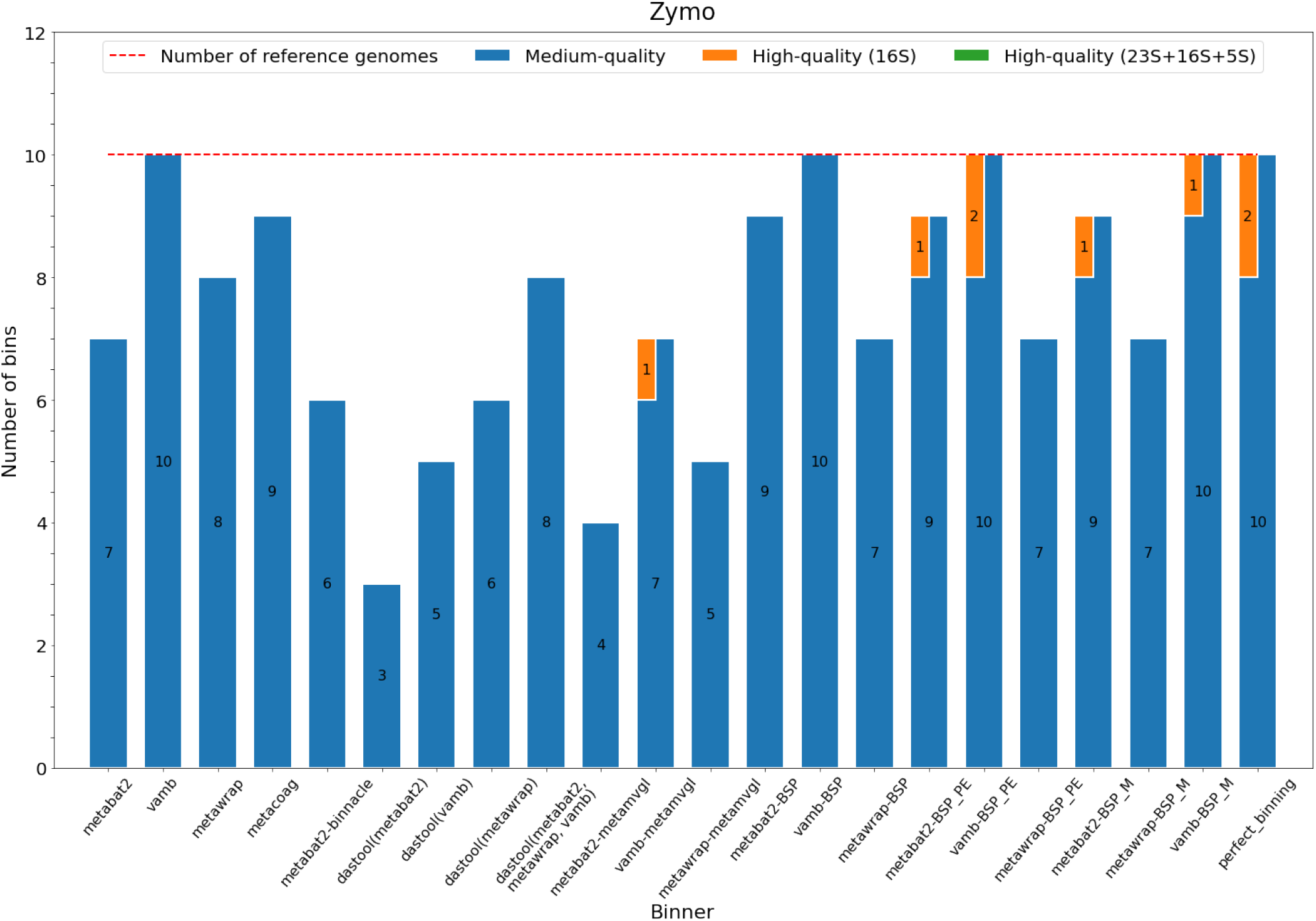
MIMAG for **Zymo** dataset. BSP denotes refining with BinSPreader in default mode, BSP_M denotes BinSPreader with multiple binning and BSP_PE denotes BinSPreader with the usage of supplementary paired-end connectivity information. perfect_binning – MAGs constructed from assembly having 100% purity and completeness. High-quality MAGs are divided into MAGs containing at least 16S/18S (orange bars) and MAGs with complete set of rRNA (green bars). Dashed line indicates number of reference genomes in dataset.

**Supplementary Figure 13:**
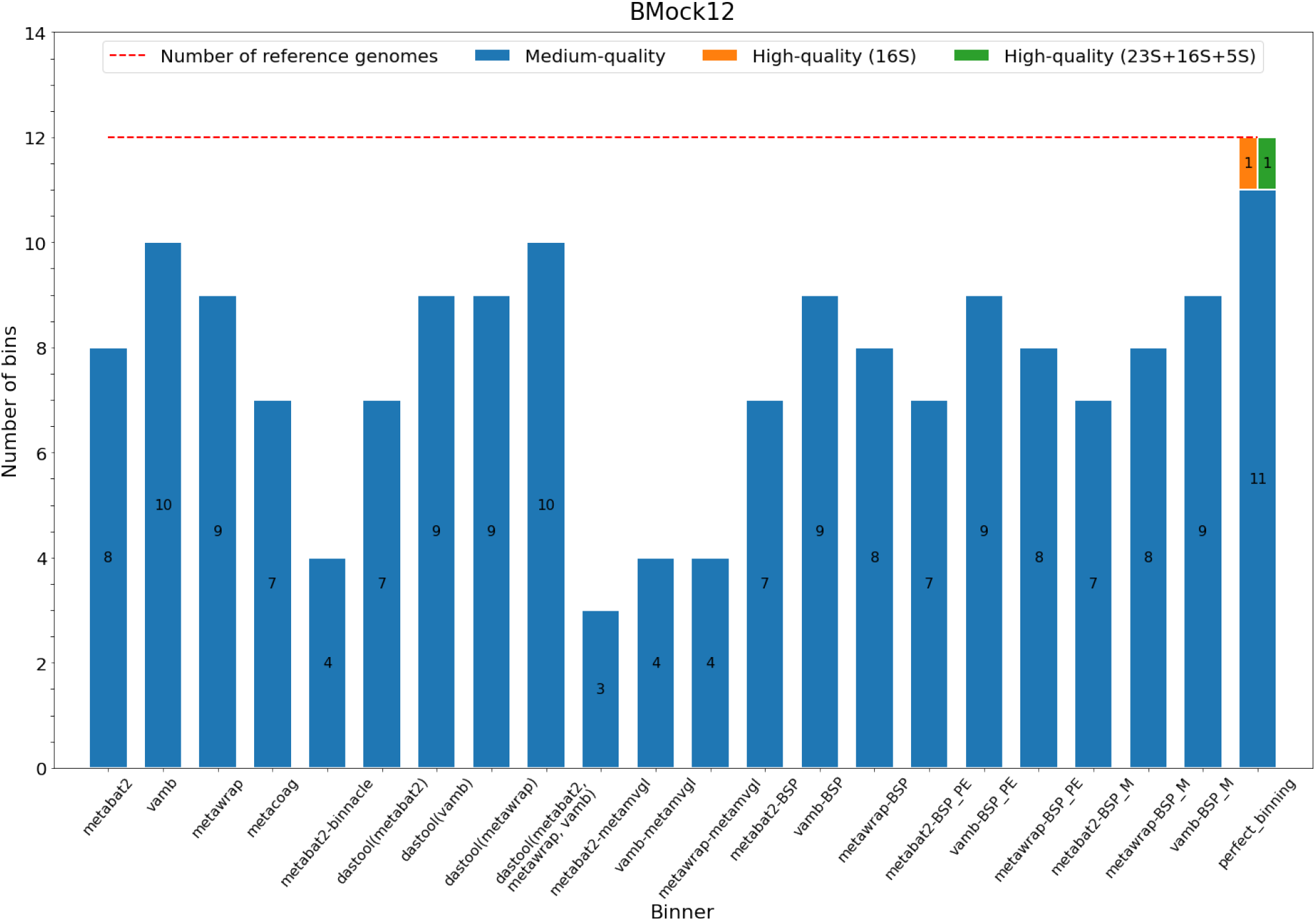
MIMAG for **BMock12** dataset. BSP denotes refining with BinSPreader in default mode, BSP_M denotes BinSPreader with multiple binning and BSP_PE denotes BinSPreader with the usage of supplementary paired-end connectivity information. perfect_binning – MAGs constructed from assembly having 100% purity and completeness. High-quality MAGs are divided into MAGs containing at least 16S/18S (orange bars) and MAGs with complete set of rRNA (green bars). Dashed line indicates number of reference genomes in dataset.

**Supplementary Figure 14:**
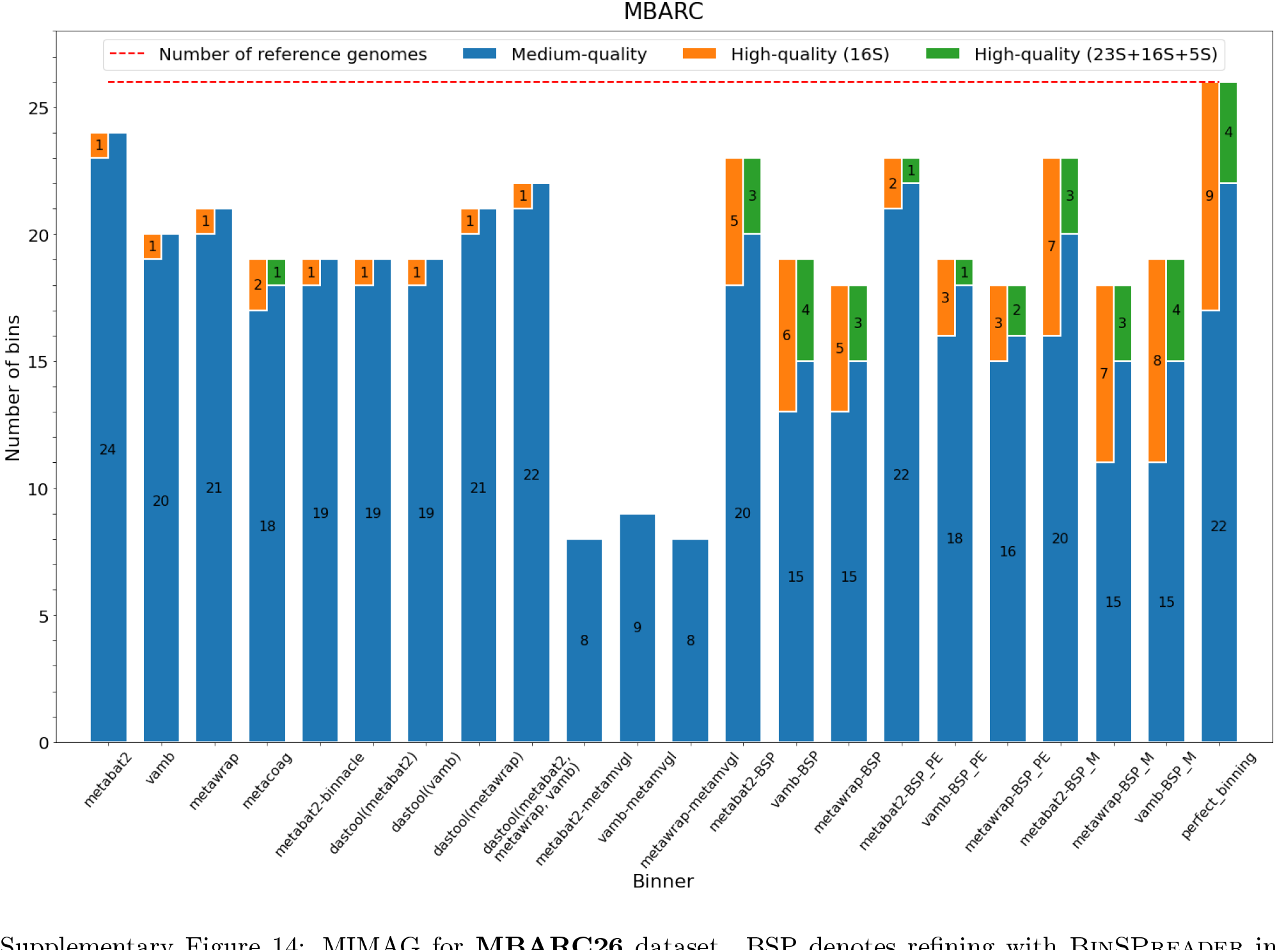
MIMAG for **MBARC26** dataset. BSP denotes refining with BinSPreader in default mode, BSP_M denotes BinSPreader with multiple binning and BSP_PE denotes BinSPreader with the usage of supplementary paired-end connectivity information. perfect_binning – MAGs constructed from assembly having 100% purity and completeness. High-quality MAGs are divided into MAGs containing at least 16S/18S (orange bars) and MAGs with complete set of rRNA (green bars). Dashed line indicates number of reference genomes in dataset.

**Supplementary Figure 15:**
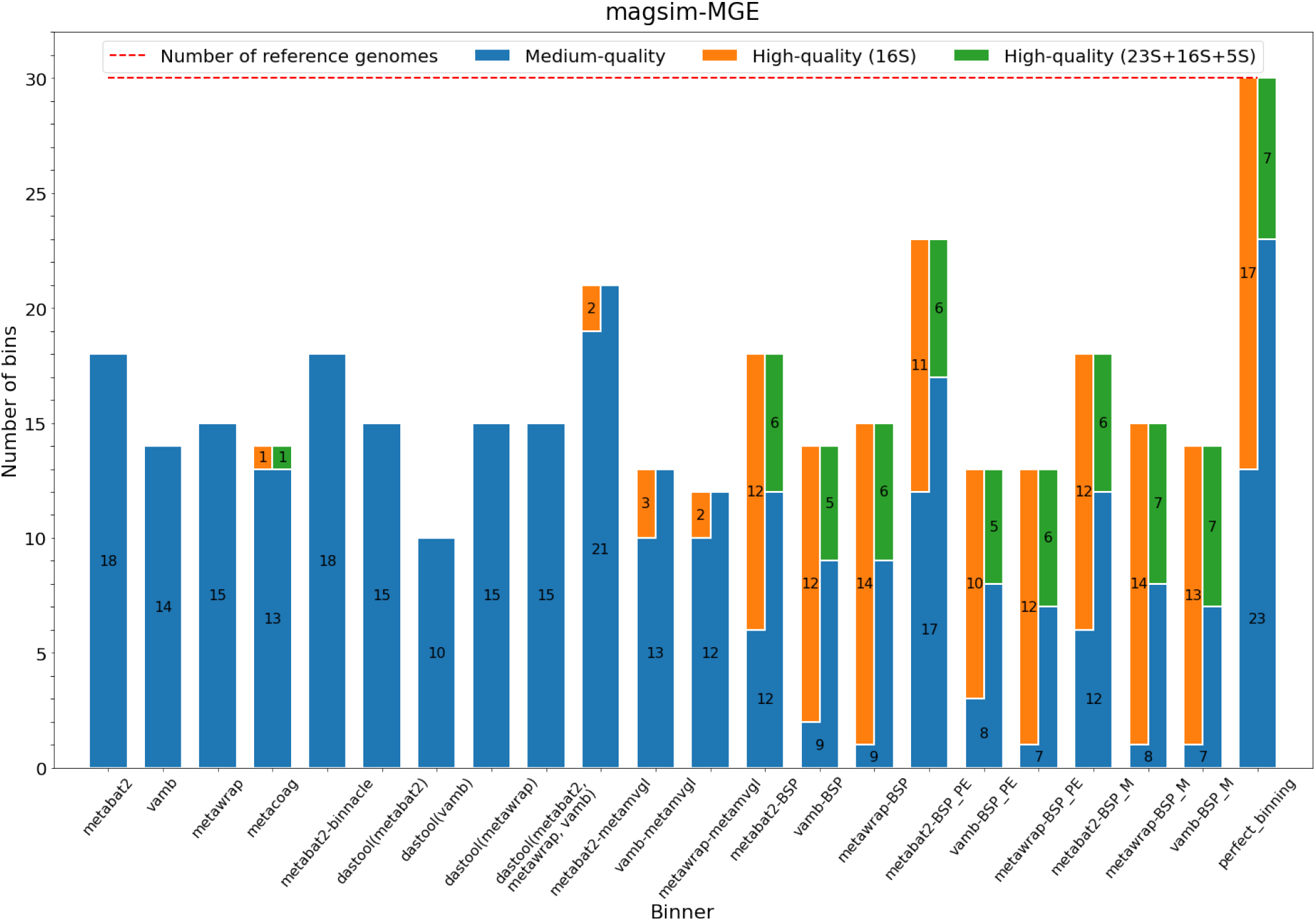
MIMAG for **magsim-MGE** dataset. BSP denotes refining with BinSPreader in default mode, BSP_M denotes BinSPreader with multiple binning and BSP_PE denotes BinSPreader with the usage of supplementary paired-end connectivity information. perfect_binning – MAGs constructed from assembly having 100% purity and completeness. High-quality MAGs are divided into MAGs containing at least 16S/18S (orange bars) and MAGs with complete set of rRNA (green bars). Dashed line indicates number of reference genomes in dataset.

**Supplementary Figure 16:**
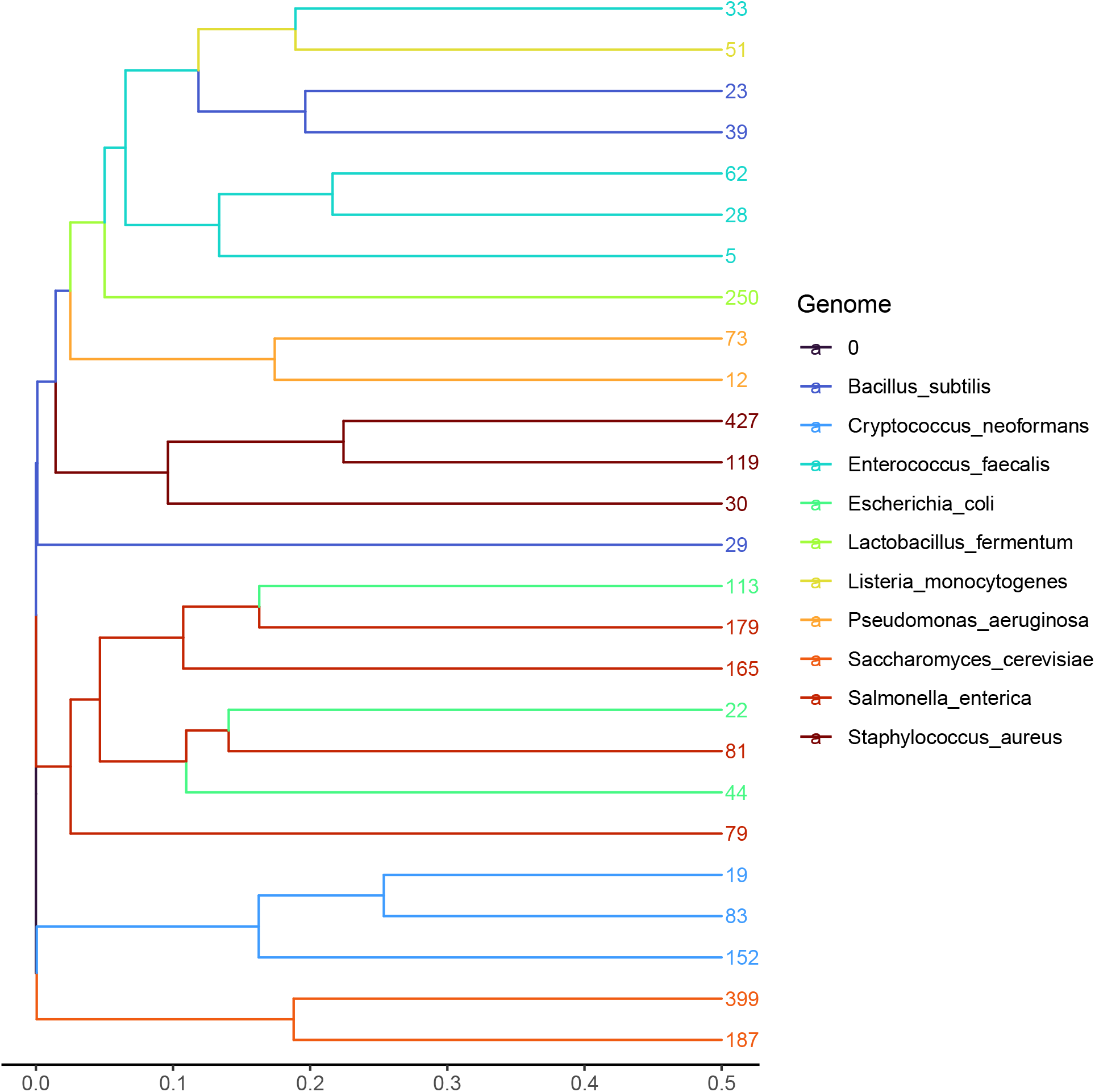
Hierarchical clustering of **Zymo** MetaBAT2 refined bins using the prob Jaccard distance between bin distributions on the assembly graph. The leafs are colored by reference and leaf numbers are bin labels. *E. coli* and *S. enterica* bins have significant overlap on the assembly graph and therefore are cross-contaminated.

**Supplementary Figure 17:**
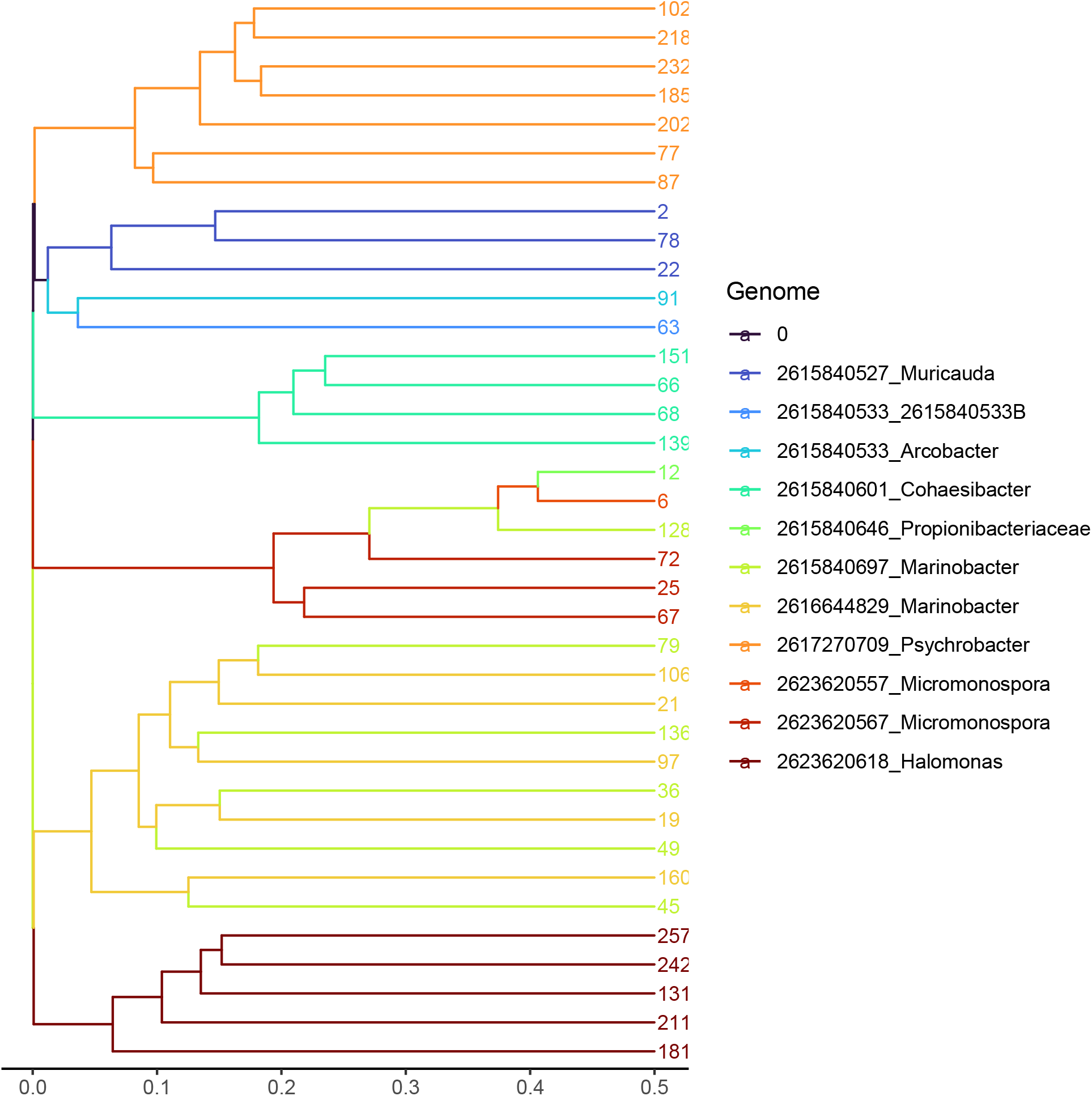
Hierarchical clustering of **BMock12** MetaBAT2 refined bins using the prob Jaccard distance between bin distributions on the assembly graph. The leafs are colored by reference and leaf numbers are bin labels. Two *Micromonospora* strains have significant overlap on the assembly graph and one of *Marinobacter* bins is clearly contaminated.

## Supplementary Tables

**Table 1:** AMBER results for **magsim-MGE** dataset. AC denotes average completeness, AP denotes average purity. The best results for each metrics are highlighted in bold.

| Tool | AC,<br>bp, % | AP,<br>bp, % | F1,<br>bp, % | % binned,<br>by length | % binned,<br>by # seq | # recovered genomes<br>depending on completeness |  |  |
| --- | --- | --- | --- | --- | --- | --- | --- | --- |
|  |  |  |  |  |  | >50% | >70% | >90% |
| Gold standard | 100.0 | 100.0 | 100.0 | 100.0 | 100.0 | 30 | 30 | 30 |
| DAS Tool + MetaBAT2 +<br>MetaWRAP + VAMB | 47.3 | **99.8** | 64.2 | 59.0 | 8.6 | 15 | **15** | 12 |
| MetaBAT2 | 69.2 | 97.2 | 80.8 | 78.5 | 30.7 | 18 | **15** | 12 |
| MetaBAT2 + BinSPreader | 72.0 | 97.0 | 82.6 | 81.0 | 35.0 | 17 | 14 | 12 |
| MetaBAT2 + BinSPreader-PE | **75.8** | 96.6 | **85.0** | **84.1** | **44.3** | **23** | 14 | 12 |
| MetaBAT2 + Binnacle | 66.0 | 97.7 | 78.8 | 78.7 | 32.5 | 18 | 12 | 9 |
| MetaBAT2 + DAS Tool | 47.1 | **99.8** | 64.0 | 58.4 | 7.9 | 15 | **15** | 12 |
| MetaBAT2 + METAMVGL | 71.7 | 95.9 | 82.0 | 77.4 | 43.9 | 21 | 13 | 9 |
| MetaCoAG | 47.3 | 97.5 | 63.7 | 57.5 | 8.6 | 12 | 12 | 10 |
| MetaWRAP | 47.4 | **99.9** | 64.3 | 59.0 | 8.6 | 15 | **15** | 13 |
| MetaWRAP + BinSPreader | 50.9 | 99.1 | 67.3 | 62.4 | 12.2 | 14 | 14 | **14** |
| MetaWRAP + BinSPreader-PE | 53.0 | 97.7 | 68.7 | 63.6 | 15.2 | 13 | 13 | 13 |
| MetaWRAP + DAS Tool | 47.4 | **99.9** | 64.3 | 59.0 | 8.6 | 15 | **15** | 13 |
| MetaWRAP + METAMVGL | 49.6 | 97.1 | 65.7 | 59.5 | 15.4 | 11 | 11 | 10 |
| VAMB | 43.5 | **99.8** | 60.6 | 52.8 | 4.6 | 14 | 13 | 8 |
| VAMB + BinSPreader | 49.6 | 98.9 | 66.1 | 59.8 | 10.1 | 14 | 13 | 13 |
| VAMB + BinSPreader-PE | 52.2 | 97.4 | 67.9 | 62.1 | 13.9 | 13 | 13 | 12 |
| VAMB + DAS Tool | 30.9 | **99.9** | 47.1 | 32.5 | 2.1 | 10 | 10 | 6 |
| VAMB + METAMVGL | 49.6 | 97.5 | 65.7 | 59.3 | 14.8 | 13 | 12 | 11 |

**Table 2:**
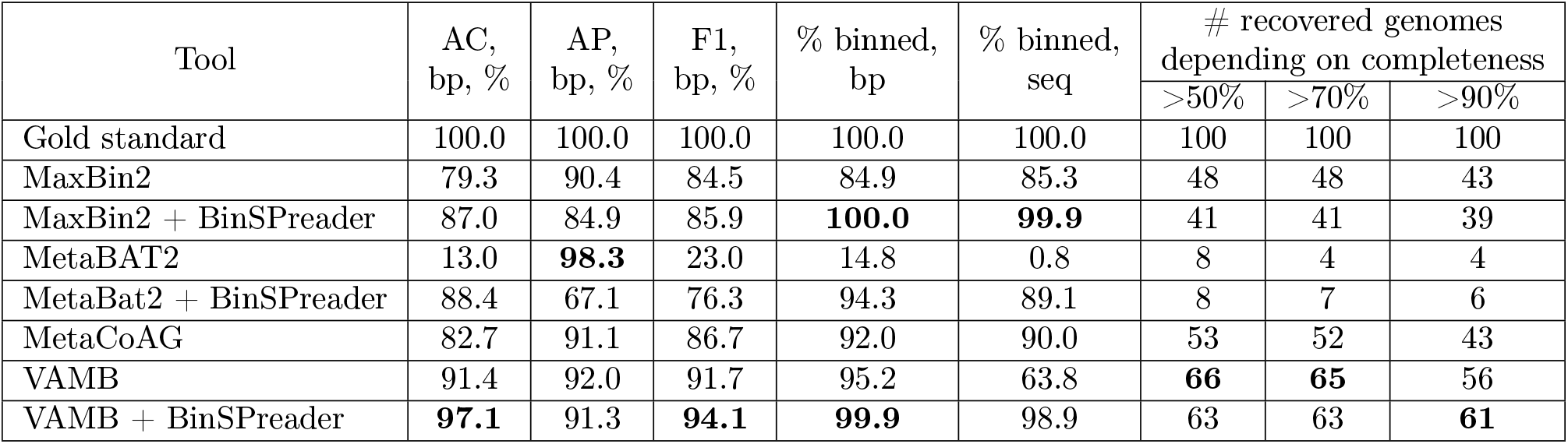
AMBER results for **simHC+** dataset. AC denotes average completeness, AP denotes average purity. The best results for each metrics are highlighted in bold.

| Tool | AC,<br>bp, % | AP,<br>bp, % | F1,<br>bp, % | % binned,<br>bp | % binned,<br>seq | # recovered genomes<br>depending on completeness |  |  |
| --- | --- | --- | --- | --- | --- | --- | --- | --- |
|  |  |  |  |  |  | >50% | >70% | >90% |
| Gold standard | 100.0 | 100.0 | 100.0 | 100.0 | 100.0 | 100 | 100 | 100 |
| MaxBin2 | 79.3 | 90.4 | 84.5 | 84.9 | 85.3 | 48 | 48 | 43 |
| MaxBin2 + BinSPreader | 87.0 | 84.9 | 85.9 | **100.0** | **99.9** | 41 | 41 | 39 |
| MetaBAT2 | 13.0 | **98.3** | 23.0 | 14.8 | 0.8 | 8 | 4 | 4 |
| MetaBat2 + BinSPreader | 88.4 | 67.1 | 76.3 | 94.3 | 89.1 | 8 | 7 | 6 |
| MetaCoAG | 82.7 | 91.1 | 86.7 | 92.0 | 90.0 | 53 | 52 | 43 |
| VAMB | 91.4 | 92.0 | 91.7 | 95.2 | 63.8 | **66** | **65** | 56 |
| VAMB + BinSPreader | **97.1** | 91.3 | **94.1** | **99.9** | 98.9 | 63 | 63 | **61** |

**Table 3:** AMBER results for mock **Zymo** dataset. AC denotes average completeness, AP denotes average purity. The best results for each metrics are highlighted in bold.

| Tool | AC,<br>bp, % | AP,<br>bp, % | F1,<br>bp, % | % binned,<br>bp | % binned,<br>seq | # recovered genomes<br>depending on completeness |  |  |
| --- | --- | --- | --- | --- | --- | --- | --- | --- |
|  |  |  |  |  |  | >50% | >70% | >90% |
| Gold standard | 100.0 | 100.0 | 100.0 | 100.0 | 100.0 | 10 | 10 | 10 |
| DAS Tool + MetaBAT2 +<br>MetaWRAP + VAMB | 77.9 | **100.0** | 87.6 | 50.6 | 26.5 | 8 | 8 | 7 |
| MetaBAT2 | 66.2 | **100.0** | 79.7 | 91.7 | 44.4 | 7 | 4 | 3 |
| MetaBAT2 + BinSPreader | 78.3 | 99.4 | 87.6 | **99.8** | 99.3 | 9 | 6 | 5 |
| MetaBAT2 + BinSPreader-PE | 79.8 | **100.0** | 88.8 | **100.0** | **100.0** | 9 | 6 | 5 |
| MetaBAT2 + Binnacle | 60.6 | **100.0** | 75.4 | 70.7 | 52.7 | 6 | 4 | 3 |
| MetaBAT2 + DAS Tool | 35.8 | **100.0** | 52.7 | 30.4 | 16.5 | 3 | 3 | 2 |
| MetaBAT2 + METAMVGL | 77.9 | 88.4 | 82.8 | 81.2 | 89.2 | 4 | 4 | 2 |
| MetaCoAG | 74.6 | 98.6 | 85.0 | 50.6 | 31.4 | 9 | 6 | 5 |
| MetaWRAP | 78.8 | **100.0** | 88.1 | 42.7 | 15.9 | 8 | 8 | 8 |
| MetaWRAP + BinSPreader | 99.7 | 95.4 | 97.5 | **99.8** | 99.4 | 7 | 7 | 7 |
| MetaWRAP + BinSPreader-PE | **99.9** | 95.1 | 97.5 | **100.0** | **100.0** | 7 | 7 | 7 |
| MetaWRAP + DAS Tool | 59.5 | **100.0** | 74.6 | 29.7 | 9.0 | 6 | 6 | 6 |
| MetaWRAP + METAMVGL | 66.5 | 94.3 | 78.0 | 80.9 | 93.3 | 5 | 3 | 1 |
| VAMB | 96.5 | **100.0** | 98.2 | 95.6 | 51.2 | **10** | **10** | **10** |
| VAMB + BinSPreader | 99.6 | **99.8** | **99.7** | **99.8** | 99.4 | **10** | **10** | **10** |
| VAMB + BinSPreader-PE | **100.0** | **100.0** | **100.0** | **100.0** | **100.0** | **10** | **10** | **10** |
| VAMB + DAS Tool | 49.1 | **100.0** | 65.9 | 23.9 | 7.7 | 5 | 5 | 5 |
| VAMB + METAMVGL | 78.0 | 94.8 | 85.6 | 86.3 | 94.5 | 7 | 5 | 3 |

**Table 4:** AMBER results for **MBARC26** dataset. AC denotes average completeness, AP denotes average purity. The best results for each metrics are highlighted in bold.

| Tool | AC,<br>bp, % | AP,<br>bp, % | F1,<br>bp, % | % binned,<br>bp | % binned,<br>seq | # recovered genomes<br>depending on completeness |  |  |
| --- | --- | --- | --- | --- | --- | --- | --- | --- |
|  |  |  |  |  |  | >50% | >70% | >90% |
| Gold standard | 100.0 | 100.0 | 100.0 | 100.0 | 100.0 | 25 | 25 | 25 |
| DAS Tool + MetaBAT2 +<br>MetaWRAP + VAMB | 88.7 | 99.2 | 93.6 | 89.2 | 50.4 | 22 | **22** | **20** |
| MetaBAT2 | 88.9 | 98.0 | 93.2 | 96.1 | 62.2 | **23** | 20 | 15 |
| MetaBAT2 + BinSPreader | 92.1 | 97.3 | 94.7 | **99.9** | 99.3 | 22 | 19 | 17 |
| MetaBAT2 + BinSPreader-PE | 91.0 | 96.2 | 93.5 | **100.0** | **99.7** | 20 | 17 | 14 |
| MetaBAT2 + Binnacle | 82.0 | 97.6 | 89.1 | 96.4 | 71.1 | 18 | 15 | 10 |
| MetaBAT2 + DAS Tool | 74.5 | 99.6 | 85.2 | 73.1 | 39.0 | 18 | 18 | 14 |
| MetaBAT2 + METAMVGL | 67.3 | 86.5 | 75.7 | 66.5 | 84.9 | 8 | 8 | 6 |
| MetaCoAG | 92.6 | 93.9 | 93.2 | 96.5 | 89.4 | 17 | 16 | 15 |
| MetaWRAP | 80.9 | **99.8** | 89.4 | 79.7 | 47.8 | 21 | 21 | 19 |
| MetaWRAP + BinSPreader | **98.9** | 92.3 | **95.5** | **99.9** | 99.3 | 18 | 18 | 17 |
| MetaWRAP + BinSPreader-PE | 96.3 | 92.3 | 94.2 | **100.0** | **99.7** | 18 | 18 | 17 |
| MetaWRAP + DAS Tool | 80.9 | **99.8** | 89.4 | 79.7 | 47.8 | 21 | 21 | 19 |
| MetaWRAP + METAMVGL | 69.5 | 90.1 | 78.5 | 69.8 | 87.4 | 8 | 8 | 5 |
| VAMB | 91.2 | 95.1 | 93.1 | 92.6 | 48.1 | 20 | 20 | 16 |
| VAMB + BinSPreader | 96.1 | 92.9 | 94.5 | **99.9** | 99.3 | 19 | 19 | 18 |
| VAMB + BinSPreader-PE | 95.3 | 92.7 | 94.0 | **100.0** | **99.7** | 19 | 19 | 17 |
| VAMB + DAS Tool | 72.8 | **99.9** | 84.2 | 71.9 | 36.1 | 19 | 19 | 16 |
| VAMB + METAMVGL | 65.8 | 89.4 | 75.8 | 67.5 | 80.4 | 9 | 8 | 6 |

**Table 5:**
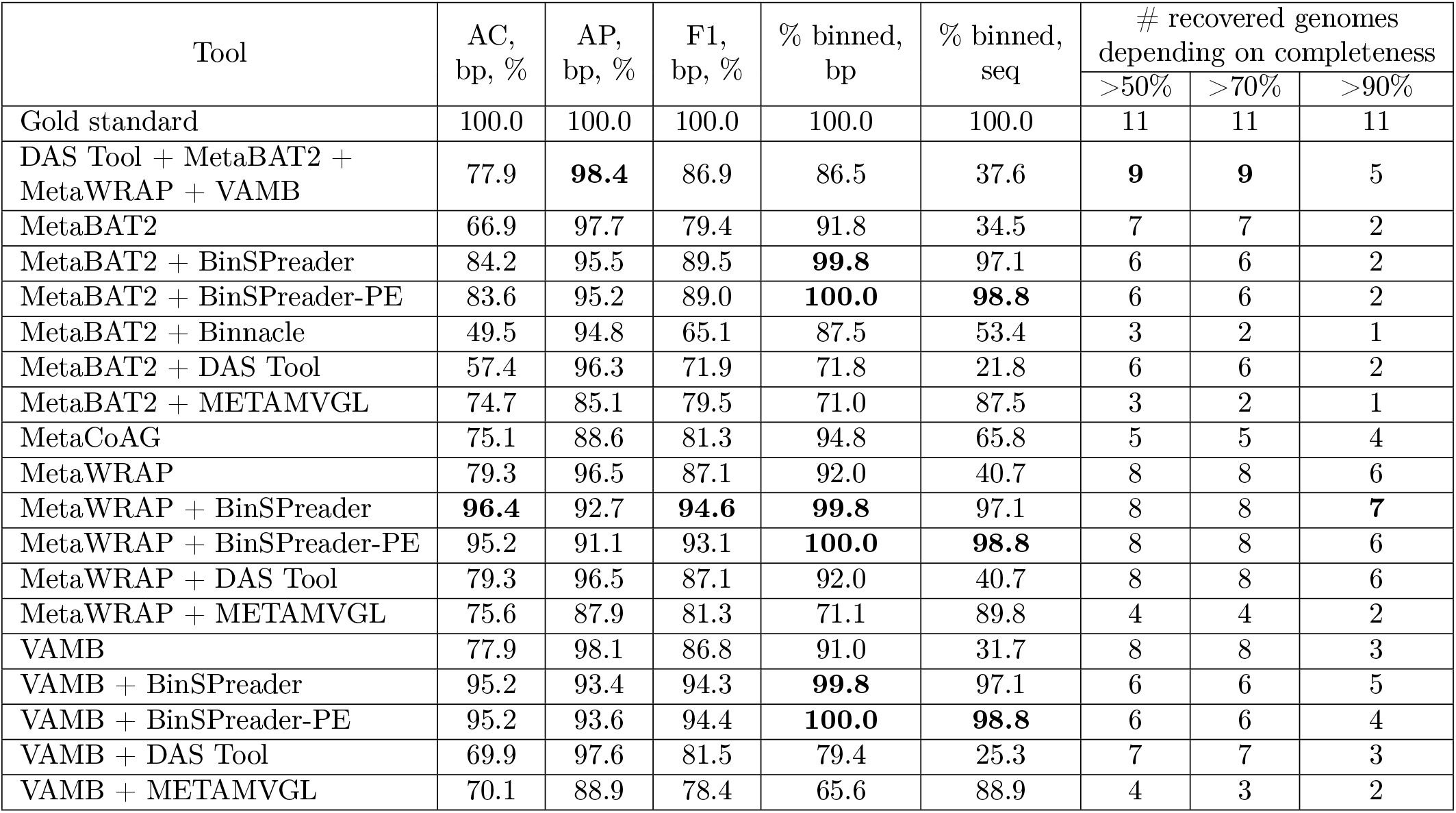
AMBER results for **BMock12** dataset. AC denotes average completeness, AP denotes average purity. The best results for each metrics are highlighted in bold.

| Tool | AC,<br>bp, % | AP,<br>bp, % | F1,<br>bp, % | % binned,<br>bp | % binned,<br>seq | # recovered genomes<br>depending on completeness |  |  |
| --- | --- | --- | --- | --- | --- | --- | --- | --- |
|  |  |  |  |  |  | >50% | >70% | >90% |
| Gold standard | 100.0 | 100.0 | 100.0 | 100.0 | 100.0 | 11 | 11 | 11 |
| DAS Tool + MetaBAT2 +<br>MetaWRAP + VAMB | 77.9 | **98.4** | 86.9 | 86.5 | 37.6 | **9** | **9** | 5 |
| MetaBAT2 | 66.9 | 97.7 | 79.4 | 91.8 | 34.5 | 7 | 7 | 2 |
| MetaBAT2 + BinSPreader | 84.2 | 95.5 | 89.5 | **99.8** | 97.1 | 6 | 6 | 2 |
| MetaBAT2 + BinSPreader-PE | 83.6 | 95.2 | 89.0 | **100.0** | **98.8** | 6 | 6 | 2 |
| MetaBAT2 + Binnacle | 49.5 | 94.8 | 65.1 | 87.5 | 53.4 | 3 | 2 | 1 |
| MetaBAT2 + DAS Tool | 57.4 | 96.3 | 71.9 | 71.8 | 21.8 | 6 | 6 | 2 |
| MetaBAT2 + METAMVGL | 74.7 | 85.1 | 79.5 | 71.0 | 87.5 | 3 | 2 | 1 |
| MetaCoAG | 75.1 | 88.6 | 81.3 | 94.8 | 65.8 | 5 | 5 | 4 |
| MetaWRAP | 79.3 | 96.5 | 87.1 | 92.0 | 40.7 | 8 | 8 | 6 |
| MetaWRAP + BinSPreader | **96.4** | 92.7 | **94.6** | **99.8** | 97.1 | 8 | 8 | **7** |
| MetaWRAP + BinSPreader-PE | 95.2 | 91.1 | 93.1 | **100.0** | **98.8** | 8 | 8 | 6 |
| MetaWRAP + DAS Tool | 79.3 | 96.5 | 87.1 | 92.0 | 40.7 | 8 | 8 | 6 |
| MetaWRAP + METAMVGL | 75.6 | 87.9 | 81.3 | 71.1 | 89.8 | 4 | 4 | 2 |
| VAMB | 77.9 | 98.1 | 86.8 | 91.0 | 31.7 | 8 | 8 | 3 |
| VAMB + BinSPreader | 95.2 | 93.4 | 94.3 | **99.8** | 97.1 | 6 | 6 | 5 |
| VAMB + BinSPreader-PE | 95.2 | 93.6 | 94.4 | **100.0** | **98.8** | 6 | 6 | 4 |
| VAMB + DAS Tool | 69.9 | 97.6 | 81.5 | 79.4 | 25.3 | 7 | 7 | 3 |
| VAMB + METAMVGL | 70.1 | 88.9 | 78.4 | 65.6 | 88.9 | 4 | 3 | 2 |

**Table 6:** AMBER results for **Zymo** dataset for bin3C binning. AC denotes average completeness, AP denotes average purity. The best results for each metrics are highlighted in bold.

| Tool | AC,<br>bp, % | AP,<br>bp, % | F1,<br>bp, % | % binned,<br>bp | % binned,<br>seq | # recovered genomes<br>depending on completeness |  |  |
| --- | --- | --- | --- | --- | --- | --- | --- | --- |
|  |  |  |  |  |  | >50% | >70% | >90% |
| Gold standard | 100.0 | 100.0 | 100.0 | 100.0 | 100.0 | 10 | 10 | 10 |
| bin3C | 96.1 | **96.7** | 96.4 | 88.3 | 59.5 | 7 | 7 | 6 |
| bin3C + BinSPreader | 99.6 | 96.5 | 98.0 | **100.0** | 99.7 | **7** | **7** | **7** |
| bin3C + BinSPreader-PE | **99.7** | 96.6 | **98.2** | **100.0** | **100.0** | **7** | **7** | **7** |

**Table 7:** CheckM results for **IC9** dataset.

| Tool | # bins | Average completeness % | Average purity % | Average F1 % |
| --- | --- | --- | --- | --- |
| MetaBAT2 | 50 | 58.5 | 99.0 | 64.7 |
| MetaBAT2 + BinSPreader | 50 | 59.8 | 97.3 | 64.9 |
| MetaBAT2 + BinSPreader-HiC | 50 | 69.1 | 92.7 | 69.6 |
| MetaBAT2 + BinSPreader-PE | 50 | 60.1 | 97.3 | 65.4 |
| MetaBAT2 + Binnacle | 57 | 51.0 | 99.2 | 57.2 |
| MetaBAT2 + DAS Tool | 23 | 87.7 | 99.4 | 92.3 |
| MetaBAT2 + METAMVGL | 46 | 46.2 | 97.1 | 53.8 |
| MetaCoAG | 37 | 82.5 | 95.4 | 86.2 |
| MetaWRAP | 31 | 94.2 | 99.4 | 96.5 |
| MetaWRAP + BinSPreader | 31 | 94.7 | 95.7 | 94.9 |
| MetaWRAP + BinSPreader-HiC | 31 | 97.3 | 88.5 | 91.1 |
| MetaWRAP + BinSPreader-PE | 31 | 94.8 | 95.0 | 94.7 |
| MetaWRAP + DAS Tool | 28 | 94.7 | 99.4 | 96.8 |
| MetaWRAP + METAMVGL | 31 | 72.1 | 95.7 | 80.2 |
| VAMB | 25 | 86.3 | 99.5 | 90.7 |
| VAMB + BinSPreader | 25 | 91.3 | 93.4 | 90.5 |
| VAMB + BinSPreader-HiC | 25 | **98.9** | 78.2 | 84.1 |
| VAMB + BinSPreader-PE | 25 | 91.4 | 91.4 | 89.4 |
| VAMB + DAS Tool | 19 | 94.3 | 99.5 | 96.5 |
| VAMB + METAMVGL | 25 | 69.2 | 93.0 | 75.6 |
| bin3C | 168 | 17.9 | **99.8** | 18.6 |
| bin3C + BinSPreader | 168 | 18.3 | 99.4 | 18.8 |
| bin3C + BinSPreader-HiC | 168 | 21.9 | 98.1 | 22.0 |
| bin3C + BinSPreader-PE | 168 | 18.4 | 99.4 | 18.8 |
| bin3C + DAS Tool | 29 | 96.4 | 99.2 | **97.7** |
| bin3C + METAMVGL | 157 | 15.1 | 99.2 | 17.0 |

**Table 8:**
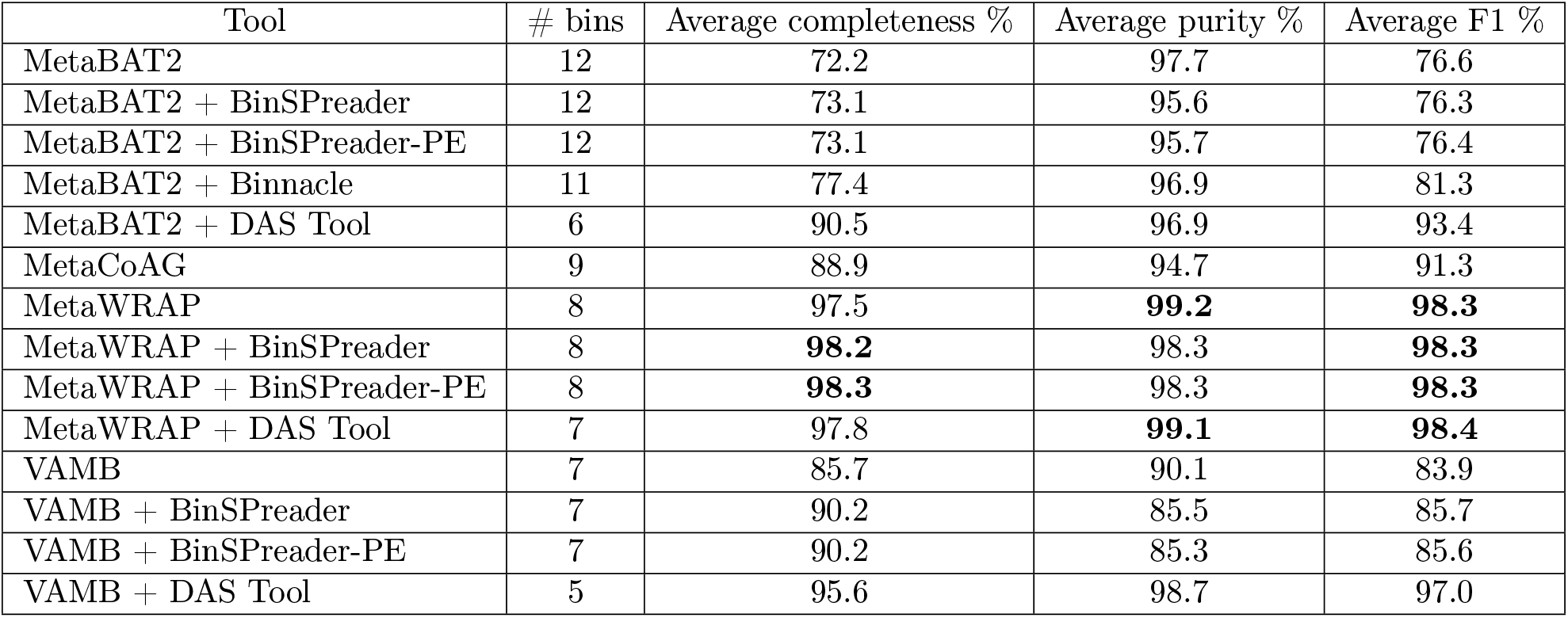
CheckM results for **Sharon** dataset.

**Table 9:** Numbers of recovered CRISPRs for mock metagenomes and **magsim-MGE** dataset. BinSPreader-PE denotes refining mode of BinSPreader utilizing additional paired-end links, and BinSPreader-Multiple denotes refining mode with multiple binning of contigs (but without paired-end data). Best results are highlighted in bold.

| Dataset/Binner | MBARC | BMock | Zymo | magsim-MGE |
| --- | --- | --- | --- | --- |
| **MetaBAT2** | 32 | 11 | 11 | 11 |
| **VAMB** | 31 | 13 | **12** | 8 |
| **metaWRAP** | 33 | 11 | **12** | 12 |
| **MetaCoAG** | 42 | 12 | **12** | 12 |
| **MetaBAT2-Binnacle** | 31 | 12 | **12** | 13 |
| **DAS TOOL (MetaBAT2)** | 26 | 5 | 5 | 11 |
| **DAS TOOL (VAMB)** | 28 | 8 | 8 | 8 |
| **DAS TOOL (metaWRAP)** | 33 | 11 | 8 | 12 |
| **DAS TOOL (MetaBAT2, metaWRAP, VAMB)** | 30 | 11 | 10 | 12 |
| **MetaBAT2-METAMVGL** | 35 | 12 | 10 | **17** |
| **VAMB-METAMVGL** | 36 | 14 | 8 | 14 |
| **metaWRAP-METAMVGL** | 36 | 14 | 6 | 16 |
| **MetaBAT2-BinSPreader** | **44** | **17** | **12** | 13 |
| **VAMB-BinSPreader** | **44** | **17** | **12** | 11 |
| **metaWRAP-BinSPreader** | **44** | **17** | **12** | 14 |
| **MetaBAT2-BinSPreader-PE** | **44** | **17** | **12** | **17** |
| **VAMB-BinSPreader-PE** | **44** | **17** | **12** | 13 |
| **metaWRAP-BinSPreader-PE** | **44** | **17** | **12** | 16 |
| **MetaBAT2-BinSPreader-Multiple** | **44** | **17** | **12** | 13 |
| **VAMB-BinSPreader-Multiple** | **44** | **17** | **12** | 11 |
| **metaWRAP-BinSPreader-Multiple** | **44** | **17** | **12** | 14 |
| **Assembly** | 44 | 17 | 12 | 23 |
| **Reference genomes** | 54 | 19 | 12 | 32 |

**Table 10:** Numbers of recovered AMR genes for mock metagenomes and **magsim-MGE** dataset. BinSPreader-PE denotes refining mode of BinSPreader utilizing paired-end links, and BinSPreader-Multiple denotes refining mode with multiple binning of contigs (but without paired-end data). Best results are highlighted in bold.

| Dataset/Binner | MBARC | BMock | Zymo | magsim-MGE |
| --- | --- | --- | --- | --- |
| MetaBAT2 | 130 | **8** | 122 | 115 |
| VAMB | 104 | **8** | 129 | 76 |
| metaWRAP | 54 | **8** | 130 | 102 |
| MetaCoAG | **138** | 6 | 133 | 106 |
| MetaBAT2-Binnacle | 135 | 7 | 129 | 117 |
| DAS TOOL (MetaBAT2) | 17 | 5 | 29 | 101 |
| DAS TOOL (VAMB) | 22 | 7 | 36 | 20 |
| DAS TOOL (metaWRAP) | 54 | **8** | 45 | 102 |
| DAS TOOL (MetaBAT2, metaWRAP, VAMB) | 55 | **8** | 81 | 102 |
| MetaBAT2-METAMVGL | 103 | 5 | 95 | 108 |
| VAMB-METAMVGL | 66 | 5 | 111 | 97 |
| metaWRAP-METAMVGL | 121 | 5 | 63 | 92 |
| MetaBAT2-BinSPreader | **139** | **8** | 133 | 135 |
| VAMB-BinSPreader | **139** | **8** | 133 | 112 |
| metaWRAP-BinSPreader | **139** | **8** | 133 | 121 |
| MetaBAT2-BinSPreader-PE | **139** | **8** | **138** | **138** |
| VAMB-BinSPreader-PE | **139** | **8** | **138** | 124 |
| metaWRAP-BinSPreader-PE | **139** | **8** | **138** | 124 |
| MetaBAT2-BinSPreader-Multiple | **139** | **8** | 133 | 135 |
| VAMB-BinSPreader-Multiple | **139** | **8** | 133 | 112 |
| metaWRAP-BinSPreader-Multiple | **139** | **8** | 133 | 121 |
| Assembly | 139 | 8 | 138 | 145 |
| Reference genomes | 153 | 11 | 182 | 220 |

**Table 11:** Numbers of recovered AMR (AntiMicrobial Resistance) genes for **IC9** dataset. BinSPreader-PE denotes refining mode of BinSPreader utilizing paired-end links, and BinSPreader-HiC denotes refining mode utilizing Hi-C links.

| **Binner** | **Genes recovered** |
| --- | --- |
| **MetaBAT2** | 70 |
| **VAMB** | 54 |
| **metaWRAP** | 70 |
| **MetaCoAG** | 134 |
| **Bin3C** | 127 |
| **MetaBAT2-Binnacle** | 73 |
| **DAS TOOL (MetaBAT2)** | 51 |
| **DAS TOOL (VAMB)** | 49 |
| **DAS TOOL (metaWRAP)** | 65 |
| **DAS TOOL (Bin3C)** | 115 |
| **MetaBAT2-METAMVGL** | 139 |
| **VAMB-METAMVGL** | 131 |
| **metaWRAP-METAMVGL** | 136 |
| **Bin3C-METAMVGL** | 146 |
| **MetaBAT2-BinSPreader** | 160 |
| **VAMB-BinSPreader** | 145 |
| **metaWRAP-BinSPreader** | 156 |
| **Bin3C-BinSPreader** | 165 |
| **MetaBAT2-BinSPreader-PE** | 161 |
| **VAMB-BinSPreader-PE** | 146 |
| **metaWRAP-BinSPreader-PE** | 157 |
| **Bin3C-BinSPreader-PE** | 166 |
| **MetaBAT2-BinSPreader-HiC** | **191** |
| **metaWRAP-BinSPreader-HiC** | **191** |
| **VAMB-BinSPreader-HiC** | **191** |
| **Bin3C-BinSPreader-HiC** | **191** |
| **Assembly** | 300 |

**Table 12:** Number of 16S/18S rRNA genes depending on their genome fraction (GF) threshold on **Zymo** dataset. The value of GF indicates the length of the assembled gene in relation to full gene. BinSPreader-PE denotes refining mode of BinSPreader utilizing paired-end links, and BinSPreader-Multiple denotes refining mode with multiple binning of contigs.

| **Zymo** |  |  |
| --- | --- | --- |
| **Binner** | **GF >50%** | **GF >90%** |
| **MetaBAT2** | 0 | 0 |
| **VAMB** | 0 | 0 |
| **metaWRAP** | 0 | 0 |
| **MetaCoAG** | 2 | 1 |
| **MetaBAT2-Binnacle** | 0 | 0 |
| **DAS TOOL (MetaBAT2)** | 0 | 0 |
| **DAS TOOL (VAMB)** | 0 | 0 |
| **DAS TOOL (metaWRAP)** | 0 | 0 |
| **DAS TOOL (MetaBAT2, metaWRAP, VAMB)** | 0 | 0 |
| **MetaBAT2-METAMVGL** | 1 | 1 |
| **VAMB-METAMVGL** | 1 | 1 |
| **metaWRAP-METAMVGL** | 2 | 1 |
| **MetaBAT2-BinSPreader** | 4 | 1 |
| **VAMB-BinSPreader** | 4 | 1 |
| **metaWRAP-BinSPreader** | 4 | 1 |
| **MetaBAT2-BinSPreader-PE** | **5** | **2** |
| **VAMB-BinSPreader-PE** | **5** | **2** |
| **metaWRAP-BinSPreader-PE** | **5** | **2** |
| **MetaBAT2-BinSPreader-Multiple** | 4 | 1 |
| **metaWRAP-BinSPreader-Multiple** | 4 | 1 |
| **VAMB-BinSPreader-Multiple** | 4 | 1 |
| **Assembly** | 5 | 2 |

**Table 13:** Number of 16S/18S rRNA genes depending on their genome fraction (GF) threshold on **magsim-MGE** dataset. The value of GF indicates the length of the assembled gene in relation to full gene. BinSPreader-PE denotes refining mode of BinSPreader utilizing paired-end links, and BinSPreader-Multiple denotes refining mode with multiple binning of contigs.

| **magsim-MGE** |  |  |
| --- | --- | --- |
| **Binner** | **GF >50%** | **GF >90%** |
| **MetaBAT2** | 1 | 1 |
| **VAMB** | 1 | 1 |
| **metaWRAP** | 2 | 2 |
| **MetaCoAG** | 7 | 6 |
| **MetaBAT2-Binnacle** | 2 | 2 |
| **DAS TOOL (MetaBAT2)** | 1 | 1 |
| **DAS TOOL (VAMB)** | 1 | 1 |
| **DAS TOOL (metaWRAP)** | 2 | 2 |
| **DAS TOOL (MetaBAT2, metaWRAP, VAMB)** | 2 | 2 |
| **MetaBAT2-METAMVGL** | 8 | 5 |
| **VAMB-METAMVGL** | 8 | 4 |
| **metaWRAP-METAMVGL** | 8 | 4 |
| **MetaBAT2-BinSPreader** | **20** | **17** |
| **VAMB-BinSPreader** | 18 | 16 |
| **metaWRAP-BinSPreader** | 18 | 16 |
| **MetaBAT2-BinSPreader-PE** | **20** | **17** |
| **VAMB-BinSPreader-PE** | 19 | **17** |
| **metaWRAP-BinSPreader-PE** | 19 | **17** |
| **MetaBAT2-BinSPreader-Multiple** | **20** | **17** |
| **metaWRAP-BinSPreader-Multiple** | 18 | 16 |
| **VAMB-BinSPreader-Multiple** | 18 | 16 |
| **Assembly** | 23 | 18 |

**Table 14:** Number of 16S/18S rRNA genes depending on their genome fraction (GF) threshold on **BMock12** dataset. The value of GF indicates the length of the assembled gene in relation to full gene. BinSPreader-PE denotes refining mode of BinSPreader utilizing paired-end links, and BinSPreader-Multiple denotes refining mode with multiple binning of contigs.

| **BMock12** |  |  |
| --- | --- | --- |
| **Binner** | **GF >50%** | **GF >90%** |
| **MetaBAT2** | 0 | 0 |
| **VAMB** | 0 | 0 |
| **metaWRAP** | 0 | 0 |
| **MetaCoAG** | 3 | 1 |
| **MetaBAT2-Binnacle** | 2 | 0 |
| **DAS TOOL (MetaBAT2)** | 0 | 0 |
| **DAS TOOL (VAMB)** | 0 | 0 |
| **DAS TOOL (metaWRAP)** | 0 | 0 |
| **DAS TOOL (MetaBAT2, metaWRAP, VAMB)** | 0 | 0 |
| **MetaBAT2-METAMVGL** | 0 | 0 |
| **VAMB-METAMVGL** | 0 | 0 |
| **metaWRAP-METAMVGL** | 0 | 0 |
| **MetaBAT2-BinSPreader** | 4 | 1 |
| **VAMB-BinSPreader** | 4 | 1 |
| **metaWRAP-BinSPreader** | 4 | 1 |
| **MetaBAT2-BinSPreader-PE** | 4 | 1 |
| **VAMB-BinSPreader-PE** | 4 | 1 |
| **metaWRAP-BinSPreader-PE** | 4 | 1 |
| **MetaBAT2-BinSPreader-Multiple** | 4 | 1 |
| **metaWRAP-BinSPreader-Multiple** | 4 | 1 |
| **VAMB-BinSPreader-Multiple** | 4 | 1 |
| **Assembly** | 4 | 1 |

**Table 15:** Number of 16S/18S rRNA genes depending on their genome fraction (GF) threshold on **MBARC26** dataset. The value of GF indicates the length of the assembled gene in relation to full gene. BinSPreader-PE denotes refining mode of BinSPreader utilizing paired-end links, and BinSPreader-Multiple denotes refining mode with multiple binning of contigs.

| **MBARC26** |  |  |
| --- | --- | --- |
| **Binner** | **GF >50%** | **GF >90%** |
| **MetaBAT2** | 2 | 1 |
| **VAMB** | 3 | 1 |
| **metaWRAP** | 3 | 1 |
| **MetaCoAG** | 12 | **9** |
| **MetaBAT2-Binnacle** | 7 | 3 |
| **DAS TOOL (MetaBAT2)** | 2 | 1 |
| **DAS TOOL (VAMB)** | 3 | 1 |
| **DAS TOOL (metaWRAP)** | 3 | 1 |
| **DAS TOOL (MetaBAT2, metaWRAP, VAMB)** | 3 | 1 |
| **MetaBAT2-METAMVGL** | 0 | 0 |
| **VAMB-METAMVGL** | 0 | 0 |
| **metaWRAP-METAMVGL** | 0 | 0 |
| **MetaBAT2-BinSPreader** | **16** | **9** |
| **VAMB-BinSPreader** | **16** | **9** |
| **metaWRAP-BinSPreader** | **16** | **9** |
| **MetaBAT2-BinSPreader-PE** | **16** | **9** |
| **VAMB-BinSPreader-PE** | **16** | **9** |
| **metaWRAP-BinSPreader-PE** | **16** | **9** |
| **MetaBAT2-BinSPreader-Multiple** | **16** | **9** |
| **metaWRAP-BinSPreader-Multiple** | **16** | **9** |
| **VAMB-BinSPreader-Multiple** | **16** | **9** |
| **Assembly** | 16 | 9 |

**Table 16:** Number of rRNA genes in bins of the **IC9** dataset depending on their genome fraction. The best results are highlighted in bold.

| Tool | 16S rRNA |  | 23S rRNA |  | 5S rRNA |  | Total rRNA |
| --- | --- | --- | --- | --- | --- | --- | --- |
|  | >50% | >80% | >50% | >80% | >50% | >80% |  |
| Assembly | 7 | 3 | 12 | 6 | 54 | 47 | 73 |
| MetaBAT2 | 0 | 0 | 0 | 0 | 7 | 6 | 7 |
| MetaBAT2 + BinSPreader | **7** | **3** | **12** | **6** | 49 | 40 | 68 |
| MetaBAT2 + BinSPreader-HiC | **7** | **3** | **12** | **6** | **52** | **44** | **71** |
| MetaBAT2 + BinSPreader-PE | **7** | **3** | **12** | **6** | 49 | 40 | 68 |
| MetaBAT2 + Binnacle | 0 | 0 | 0 | 0 | 7 | 7 | 7 |
| MetaBAT2 + DAS Tool | 0 | 0 | 0 | 0 | 5 | 4 | 5 |
| MetaBAT2 + METAMVGL | 1 | 0 | 3 | 1 | 28 | 23 | 32 |
| MetaCoAG | 5 | **3** | 10 | 5 | 21 | 19 | 36 |
| MetaWRAP | 0 | 0 | 0 | 0 | 8 | 8 | 8 |
| MetaWRAP + BinSPreader | **7** | **3** | **12** | **6** | 48 | 40 | 67 |
| MetaWRAP + BinSPreader-HiC | **7** | **3** | **12** | **6** | **52** | **44** | **71** |
| MetaWRAP + BinSPreader-PE | **7** | **3** | **12** | **6** | 48 | 40 | 67 |
| MetaWRAP + DAS Tool | 0 | 0 | 0 | 0 | 8 | 8 | 8 |
| MetaWRAP + METAMVGL | 0 | 0 | 3 | 1 | 26 | 21 | 29 |
| VAMB | 1 | 1 | 5 | 3 | 6 | 6 | 12 |
| VAMB + BinSPreader | **7** | **3** | 11 | 5 | 47 | 39 | 65 |
| VAMB + BinSPreader-HiC | **7** | **3** | **12** | **6** | 51 | **44** | 70 |
| VAMB + BinSPreader-PE | **7** | **3** | **12** | **6** | 47 | 40 | 66 |
| VAMB + DAS Tool | 1 | 1 | 5 | 3 | 5 | 5 | 11 |
| VAMB + METAMVGL | 0 | 0 | 3 | 0 | 22 | 18 | 25 |
| bin3C | 6 | **3** | **12** | **6** | 18 | 16 | 36 |
| bin3C + BinSPreader | **7** | **3** | **12** | **6** | 49 | 40 | 68 |
| bin3C + BinSPreader-HiC | **7** | **3** | **12** | **6** | **52** | **44** | **71** |
| bin3C + BinSPreader-PE | **7** | **3** | **12** | **6** | 49 | 40 | 68 |
| bin3C + DAS Tool | 5 | 2 | 8 | 4 | 15 | 14 | 28 |
| bin3C + METAMVGL | 0 | 0 | 3 | 1 | 21 | 17 | 24 |

**Table 17:** Number of recovered rRNA genes in bins of the **Sharon** dataset depending on their genome fraction. The best results for each metrics are highlighted in bold.

| Tool | 16S rRNA |  | 23S rRNA |  | 5S rRNA |  | Total rRNA |
| --- | --- | --- | --- | --- | --- | --- | --- |
|  | >50% | >90% | >50% | >90% | >50% | >90% |  |
| Assembly | 7 | 1 | 6 | 4 | 16 | 14 | 29 |
| MetaBAT2 | 0 | 0 | 0 | 0 | 1 | 1 | 1 |
| MetaBAT2 + BinSPreader | 3 | **1** | 3 | **3** | **12** | **10** | 18 |
| MetaBAT2 + BinSPreader-PE | **4** | **1** | **4** | **3** | **12** | **10** | **20** |
| MetaBAT2 + Binnacle | 1 | 0 | 0 | 0 | 1 | 1 | 2 |
| MetaBAT2 + DAS Tool | 0 | 0 | 0 | 0 | 1 | 1 | 1 |
| MetaCoAG | 3 | 0 | 1 | 1 | 2 | 2 | 6 |
| MetaWRAP | 1 | 0 | 0 | 0 | 1 | 1 | 2 |
| MetaWRAP + BinSPreader | 3 | **1** | 3 | **3** | 8 | 6 | 14 |
| MetaWRAP + BinSPreader-PE | **4** | **1** | **4** | **3** | **12** | **10** | **20** |
| VAMB | 1 | **1** | 1 | 1 | 2 | 2 | 4 |
| VAMB + BinSPreader | 3 | **1** | 3 | **3** | 8 | 6 | 14 |
| VAMB + BinSPreader-PE | **4** | **1** | **4** | **3** | **12** | **10** | **20** |
| VAMB + DAS Tool | 1 | **1** | 1 | 1 | 2 | 2 | 4 |

## Notes

https://figshare.com/projects/BinSPreader/132425

https://cab.spbu.ru/software/binspreader/

### Competing Interest Statement

The authors have declared no competing interest.

